# Caspase-2 is essential for proliferation and self-renewal of nucleophosmin-mutated acute myeloid leukemia

**DOI:** 10.1101/2023.05.29.542723

**Authors:** Dharaniya Sakthivel, Alexandra N. Brown-Suedel, Francesca Keane, Shixia Huang, Kenneth Mc Sherry, Chloé I. Charendoff, Kevin P. Dunne, Dexter J. Robichaux, BaoChau Le, Crystal S. Shin, Alexandre F. Carisey, Jonathan M. Flanagan, Lisa Bouchier-Hayes

**Affiliations:** Department of Pediatrics, Division of Hematology-Oncology, Baylor College of Medicine, Houston, TX 77030, USA; Texas Children's Hospital William T. Shearer Center for Human Immunobiology, Houston, TX, 77030, USA; Department of Molecular and Human Genetics, Baylor College of Medicine, Houston, TX 77030, USA; Advanced Technology Cores, Department of Molecular and Cellular Biology, Department of Education, Innovation & Technology, Dan L. Duncan Cancer Center, Baylor College of Medicine, Houston, TX 77030, USA; Michael E. DeBakey Department of Surgery, Baylor College of Medicine, Houston, TX 77030 USA; Department of Molecular and Cellular Biology, Baylor College of Medicine, Houston, TX, 77030, USA

**Keywords:** Acute myeloid leukemia, nucleophosmin, caspase-2, NPM1c+, nucleolus

## Abstract

Mutation in nucleophosmin (NPM1) causes relocalization of this normally nucleolar protein to the cytoplasm (*NPM1c+*). Despite NPM1 mutation being the most common driver mutation in cytogenetically normal adult acute myeloid leukemia (AML), the mechanisms of NPM1c+-induced leukemogenesis remain unclear. Caspase-2 is a pro-apoptotic protein activated by NPM1 in the nucleolus. Here, we show that caspase-2 is also activated by NPM1c+ in the cytoplasm, and DNA damage-induced apoptosis is caspase-2-dependent in *NPM1c+* AML but not in *NPM1wt* cells. Strikingly, in *NPM1c+* cells, loss of caspase-2 results in profound cell cycle arrest, differentiation, and down-regulation of stem cell pathways that regulate pluripotency including impairment in the AKT/mTORC1 and Wnt signaling pathways. In contrast, there were minimal differences in proliferation, differentiation, or the transcriptional profile of *NPM1wt* cells with and without caspase-2. Together, these results show that caspase-2 is essential for proliferation and self-renewal of AML cells that have mutated NPM1. This study demonstrates that caspase-2 is a major effector of NPM1c+ function and may even be a druggable target to treat *NPM1c+* AML and prevent relapse.

## Introduction

Accounting for 80% of adult acute leukemia, acute myeloid leukemia (AML) is an aggressive malignant blood cancer that impairs hematopoiesis resulting in the accumulation of immature blood cells (Noone AM, 2018). Among the known driver mutations for AML, mutation of nucleophosmin (NPM1) is the most common, occurring in 60% of adult cytogenetically normal AML (Falini et al., 2005). Nucleophosmin is a ubiquitous, multi-functional phosphoprotein that localizes in the nucleolus and primarily functions in ribosomal biogenesis (Grisendi et al., 2006). The most common mutation of NPM1 (Type A) results from a four base pair insertion in the C-terminus that harbors a nucleolar localization signal (NuLS) (Falini et al., 2005). This insertion disrupts the NuLS and instead generates a leucine-rich strong nuclear export motif (NES) resulting in constitutive nuclear export of NPM1. Thus, the mutant version of NPM1 is referred to as NPM1 cytoplasmic positive or *NPM1c+* (Falini et al., 2005).

Although mutation in NPM1 is a leukemia-generating event, *NPM1c+* AML has a favorable prognostic index compared to *NPM1wt* AML. As such, it has been defined as a distinct leukemia entity by the World Health Organization (Arber et al., 2022). Compared to *NPM1wt* AML, *NPM1c+* AML is associated with high rates of complete molecular remission following standard induction therapy (Falini et al., 2005). The improved response to treatment is thought to be a result of increased sensitivity of NPM1c+ cells to apoptosis induced by chemotherapeutic drugs (Martelli et al., 2015). However, the mechanisms by which NPM1c+ can both drive leukemogenesis and improve treatment responses are unclear.

We previously reported that NPM1 is an upstream activator of the pro-apoptotic protein caspase-2 in the nucleolus (Ando et al., 2017). Caspase-2 is an initiator caspase that is activated by proximity-induced dimerization of inactive monomers following recruitment to specific high molecular weight protein complexes known as activation platforms (Brown-Suedel and Bouchier-Hayes, 2020). The activation platform for caspase-2 is the PIDDosome, comprising of PIDD and RAIDD (Tinel and Tschopp, 2004). In response to DNA damage, we showed that NPM1 provides a scaffold for PIDDosome formation and caspase-2 activation in the nucleolus (Ando et al., 2017). Cytoplasmic activation of caspase-2 did not require PIDD or NPM1, indicating that two separate activation platforms are assembled in response to DNA damage: one in the cytoplasm and one in the nucleolus. The two different subcellular localizations of caspase-2 activation suggest that, depending on where it is activated in the cell, caspase-2 may have access to different substrates leading to distinct downstream functions. Therefore, it is possible that the subcellular localization of caspase-2 activation is a major determinant of its downstream function, and caspase-2 may even regulate non-apoptotic functions from the nucleolus in an NPM1-dependent manner. Because NPM1c+ changes the localization of NPM1, this could have a dramatic effect on the downstream activation of caspase-2 and resulting functional outcomes.

While caspase-2 is considered a pro-apoptotic protein, it has been shown to have non-apoptotic roles. For example, caspase-2-deficient cells proliferate at higher rates and have increased replication stress (Boice et al., 2022; Ho et al., 2009). As a result, caspase-2-deficient cells and tissues show many features of persistent DNA damage and genomic instability (Dorstyn et al., 2012; Parsons et al., 2013; Puccini et al., 2013). In caspase-2-deficient murine models, this genomic instability manifests as accelerated aging or, in certain tumor models, accelerated tumorigenesis (Ho et al., 2009; Parsons et al., 2013; Puccini et al., 2013; Terry et al., 2014; Zhang et al., 2007). In aged mice, loss of caspase-2 leads to increased DNA damage in hematopoietic stem cells and increased hematopoiesis (Dawar et al., 2016). Despite the fact that caspase-2 is a proven tumor suppressor in the Eµ-Myc lymphoma mouse model, and increased caspase-2 expression is associated with somewhat improved survival and drug sensitivity in AML (Dawar et al., 2016; Holleman et al., 2005), a role for caspase-2 in regulating hematopoiesis in leukemia has not been studied.

*NPM1c+* AML is characterized by an increased stem cell molecular signature (Alcalay et al., 2005). *NPM1c+* is associated with increased expression of several homeobox HOX genes, which are highly expressed in hematopoietic stem cells (Alcalay et al., 2005). This HOX gene signature is specific to *NPM1c+* AML and has been proposed to be important for leukemia development (Eklund, 2011). This is supported by studies showing that enforced relocalization of NPM1c+ back to the nucleus results in the downregulation of the HOX/MEIS1 network, cell cycle arrest, and terminal differentiation of AML cells (Brunetti et al., 2018).

Given the ability of NPM1 to activate caspase-2, and proposed roles for caspase-2 in cell cycle regulation and hematopoiesis, we set out to determine the downstream consequences of NPM1- dependent caspase-2 activation in *NPM1wt* and *NPM1c+* AML cells. Our initial prediction was that loss of caspase-2 would increase proliferation and/or enhance resistance to apoptosis in one or both of these cell types. Instead, we found minimal effects of caspase-2 loss in *NPM1wt* cells. In contrast, in *NPM1c+* cells, loss of caspase-2 reduced apoptosis. Loss of caspase-2 also had profound effects on cell division in *NPM1c+* cells, but these were opposite to our initial hypothesis that blocking caspase-2 would increase proliferation. Rather, we made the surprising discovery that, in *NPM1c+* AML, caspase-2 is essential for proliferation and self-renewal.

## Results

### Loss of caspase-2 inhibits apoptosis when NPM1 is localized to the cytoplasm

To investigate how mutation in NPM1 and the subsequent localization changes affect cellular function, we used OCI AML-2 and OCI AML-3 cells. These cells express *NPM1wt* (nucleolus) and *NPM1c+* (cytoplasm), respectively (Figure 1A). For simplicity, we will refer to the OCI-AML2 cells as *NPM1wt* and the OCI-AML3 cells as *NPM1c+* throughout this manuscript. To demonstrate that NPM1c+ increases sensitivity to apoptosis in AML cells, we incubated *NPM1wt* and *NPM1c+* cells with the DNA-damaging agents, doxorubicin, daunorubicin, and etoposide (Figure 1B–D). These drugs are commonly used in AML chemotherapy. In each case, the *NPM1c+* cells were significantly more sensitive to apoptosis over a range of concentrations.

**Figure 1:**
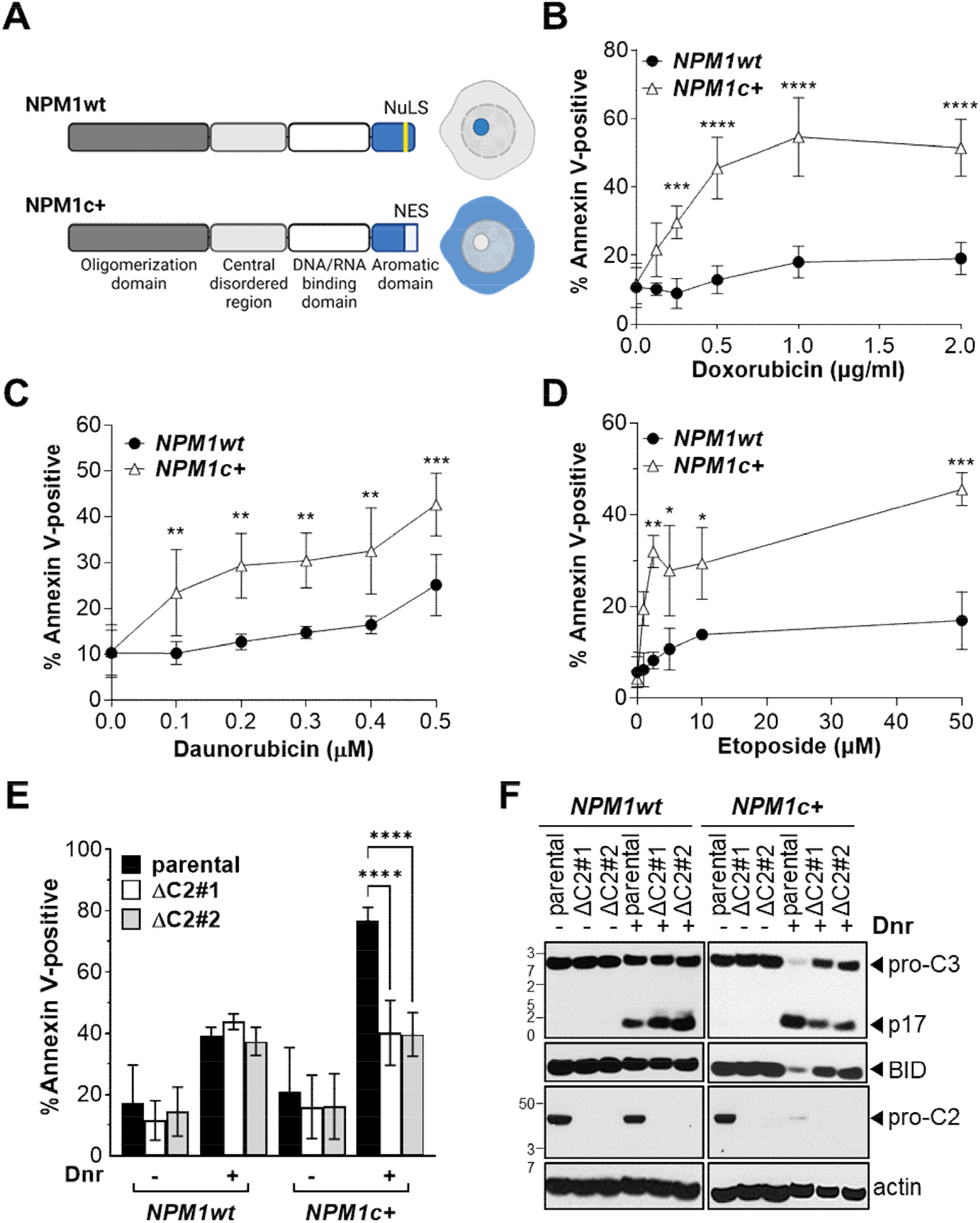
Increased sensitivity of NPM1c+ AML cells to DNA damage-induced apoptosis is caspase-2-dependent. (A) Schematic representation showing the protein domains of NPM1 and NPM1c+. The nucleolar localization signal (NuLS) sequence is indicated in NPM1, which is mutated to a nuclear exclusion signal (NES) in NPM1c+. The localization of NPM1 in *NPM1wt* cells (OCI-AML-2) and in *NPM1c+* cells (OCI-AML-3) is shown in the nucleolus or cytoplasm, respectively, in blue. (B–D) *NPM1wt* and *NPM1c+* were treated with the indicated doses of doxorubicin (B), daunorubicin (C), and etoposide (D) for 16 h. Apoptosis was assessed by flow cytometry for Annexin V binding. Results are the mean of three to four independent experiments plus or minus standard deviation *p < 0.05, **p < 0.01, ***p < 0.001, ****p < 0.0001. (E) *NPM1wt* and *NPM1c+* cells and their respective CRISPR/Cas9-generated caspase-2-deficient clones (ΔC2) were treated with and without daunorubicin (Dnr, 0.5 µM) for 16 h. The percentage of cells undergoing apoptosis was measured by flow cytometry for Annexin V binding. Results are the mean of four independent experiments plus or minus standard deviation. ****p < 0.0001. (F) Parental *NPM1wt*, *NPM1c+* cells, and the ΔC2 clones of each were treated overnight with daunorubicin (0.5 µM). Lysates were immunoblotted for caspase-3, BID, and caspase-2. Pro-C3 and pro-C2 = the pro-form of caspase-3 and caspase-2; p17 = the large catalytic subunit. Actin was used as a loading control. Results are representative of three independent experiments.

To examine the role of caspase-2 in apoptosis in *NPM1wt* and *NPM1c+* cells, we used CRISPR/Cas9 to generate caspase-2-deficient *NPM1wt* and *NPM1c+* cells. We identified two single-cell clones of each line that showed complete deletion of caspase-2. Following exposure to daunorubicin, loss of caspase-2 in *NPM1wt* cells had little effect on the susceptibility of the cells to apoptosis (Figure 1E). Strikingly, in *NPM1c+* cells, loss of caspase-2 resulted in significantly decreased apoptosis compared to the parental line (Figure 1E). Interestingly, apoptosis of *NPM1c+* cells was not completely blocked in the absence of caspase-2; rather it was decreased to the level induced in the *NPM1wt* cells. This suggests that the increased sensitivity of *NPM1c+* cells to apoptosis compared to *NPM1wt* cells is caspase-2 dependent.

Caspase-2 induces apoptosis through BID cleavage, which results in mitochondrial outer membrane permeabilization (MOMP) and downstream caspase-3 activation (Bonzon et al., 2006; Upton et al., 2008). To determine the apoptotic pathway induced by caspase-2 in *NPM1wt* and *NPM1c+* AML cells, we measured the protein levels of BID and caspase-3. In *NPM1wt* parental cells, we observed caspase-3 cleavage following daunorubicin treatment. Loss of caspase-2 resulted in a slight increase in the amount of the cleaved p17 caspase-3 subunit after daunorubicin treatment (Figure 1F). Consistent with the higher rates of apoptosis in *NPM1c+* parental cells, caspase-3 cleavage was increased compared to the *NPM1wt* cells. We also observed the disappearance of the full-length caspase-2 band in *NPM1c+* cells with treatment, suggesting caspase-2 activation. In the absence of caspase-2, the level of caspase-3 cleavage was decreased, as measured by lower levels of the p17 subunit and higher levels of the full-length caspase. Caspase-2 deficiency in *NPM1c+* cells lead to a reduction of caspase-3 cleavage to the level of that observed in the *NPM1wt* cells. We were unable to detect the cleaved form of BID in these experiments; therefore, we used the level of intact BID as an indicator of cleavage. In *NPM1c+* cells, full-length BID was reduced following daunorubicin treatment, while there was a minimal reduction of full-length BID in *NPM1wt* cells. Deletion of caspase-2 restored the levels of BID in daunorubicin-treated *NPM1c+* cells. Together, these results suggest that the increased sensitivity to apoptosis in *NPM1c+* cells is the result of caspase-2-induced BID cleavage and downstream caspase-3 activation.

### Caspase-2 is activated in the same cellular compartment as NPM1

We previously reported that caspase-2 is activated in the nucleolus by NPM1-mediated PIDDosome assembly (Ando et al., 2017). We showed that NPM1 binds to PIDD through its central disordered region (Ando et al., 2017). Since the NPM1c+ mutation is in the C-terminus of NPM1, this would suggest that NPM1c+ retains the ability to activate caspase-2. Despite this, we showed that caspase-2 cleavage was impaired in *NPM1c+* cells treated with the combination of irradiation and a Chk1 inhibitor (Ando et al., 2017). This finding is not consistent with the data shown in Figure 1E, which shows reduced apoptosis in caspase-2-deficient *NPM1c+* cells and, hence, suggests caspase-2 activation in *NPM1c+* cells. Given that caspase-2 cleavage is not the most reliable indicator of activation (Baliga et al., 2004), we decided to revisit the question of whether NPM1c+ can activate caspase-2 in the cytoplasm. We first determined the sub-cellular localization of caspase-2 in both cell types. We fractionated *NPM1wt* and *NPM1c+* cells into cytoplasmic, nuclear, and nucleolar fractions (Figure 2A). The purity of the nucleolar fraction was confirmed by probing for the nucleolar protein fibrillarin. In *NPM1wt* cells, NPM1 was detected in both the nucleolus and cytoplasm. This finding reflects the normal nucleus-cytoplasm shuttling activity of NPM1 (Borer et al., 1989). Caspase-2 was concentrated in the nucleolus of these *NPM1wt* cells. As expected, in *NPM1c+* cells, NPM1 was concentrated in the cytoplasmic and nuclear fractions but not in the nucleolus. Caspase-2 was also detected in the cytoplasmic and nuclear fractions but not in the nucleolus of *NPM1c+* cells. This suggests that NPM1 and caspase-2 are localized in the same sub-cellular fractions in *NPM1*wt and *NPM1c+* AML cells (Figure 2A).

**Figure 2:**
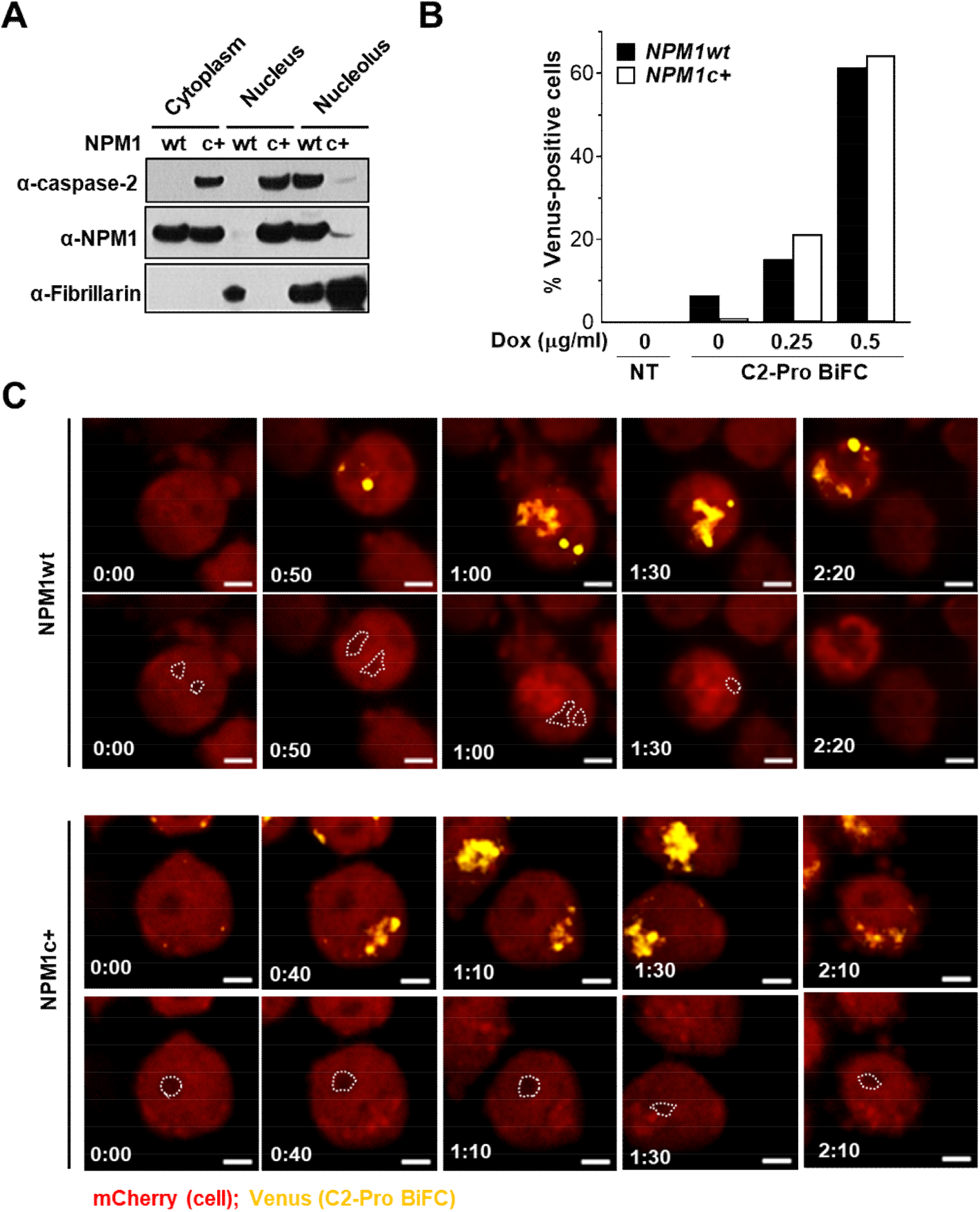
**Caspase-2 is activated in the same cellular compartment as NPM1.** (A) OCI-AML-2 (*NPM1wt*) and OCI-AML-3 (*NPM1c+*) cells were fractionated into the cytosol, nucleoplasm, and nucleolus and immunoblotted for caspase-2, NPM1, and fibrillarin (nucleolus). (B) *NPM1wt* and *NPM1c+* cells stably expressing C2-Pro VC-2A-C2-Pro VN-2A-mCherry (C2-Pro BiFC) were treated with the indicated doses of doxorubicin for 16 h in the presence of the caspase inhibitor qVD-OPH (5 µM) to prevent cell death caused by apoptosis. Cells were assessed for the percentage of Venus-positive cells with and without treatment using an imaging flow cytometer to determine caspase-2 activation at the single-cell level. (C) *NPM1wt* and *NPM1c+* C2-Pro BiFC cells treated with daunorubicin (0.5 µM) were imaged by confocal microscopy for 16 h. Representative confocal time-lapse images show cells (*red*) and caspase-2 BiFC (*yellow*) in the nucleolus (*dashed outline*) or cytoplasm (at the periphery). Bar, 10 µm.

To determine the localization of caspase-2 activation in *NPM1wt* and *NPM1c+* cells, we used caspase-2 bimolecular fluorescence complementation (BiFC). BiFC uses non-fluorescent fragments of the yellow fluorescent protein Venus that can associate to reform the fluorescent complex when fused to interacting proteins (Shyu et al., 2006). When the pro-domain of caspase-2 (C2-Pro) is fused to each half of split Venus (VC or VN), the recruitment of caspase-2 to its activation platform and the subsequent induced proximity results in enforced association of the two Venus halves. Thus, Venus fluorescence acts as a read-out for caspase-2 induced proximity, the proximal step in its activation (Bouchier-Hayes et al., 2009). We previously described a lentiviral bicistronic construct where the C2 Pro-VC and C2 Pro-VN are expressed in a single vector separated by the viral 2A self-cleaving peptide that is also linked to an mCherry reporter of expression (Ando et al., 2017). We used this construct to create OCI-AML2 and OCI-AML3 cell lines stably expressing the caspase-2 BiFC reporter.

We treated the NPM1wt and NPM1c+ C2-Pro BiFC cells with doxorubicin or daunorubicin. Following treatment, caspase-2 was activated, as measured by Venus fluorescence (Figure 2 B, C, Supplemental Figure S1). The overall activation of caspase-2 was the same for both *NPM1wt* and *NPM1c+* cells (Figure 2B). An advantage of these cells is that mCherry fluorescence is excluded from the nucleolus allowing us to precisely assess nucleolar localization of the BiFC. Using imaging flow cytometry, we observed that, in *NPM1wt* cells, caspase-2 BiFC was detected in the nucleolus while, in *NPM1c+* cells, caspase-2 BiFC was only detected outside of the nucleolus at the periphery of the cells (Supplemental Figure S1A). We confirmed these localizations using time-lapse confocal microscopy (Figure 2C, Supplemental Movie S1, S2). Because AML cells are suspension cells, they are difficult to track by imaging. To circumvent this problem, we plated them in a custom-designed micro-scaffold to keep them restricted to small square-shaped microwells with an area of 50 µm^2^ (Supplemental Figure 1B). Thus, we were able to to track the cells over time by microscopy. The caspase-2 BiFC appeared with similar kinetics in both cell types and reached a similar overall level of brightness. In the *NPM1wt* cells, caspase-2 BiFC began in the nucleous and, with time, started to appear in additional regions of the cell. In contrast, in *NPM1c+* cells, caspase-2 BiFC was excluded from the nucleolus throughout the time-lapse. Together, these results suggest that caspase-2 is activated in different compartments of the cell in a manner that is dictated by the sub-cellular localization of NPM1. The distinct localization pattern of caspase-2 activation may explain the different caspase-2 dependencies of apoptosis in *NPM1wt* and *NPM1c+* cells.

### Loss of caspase-2 differentially impacts the cell cycle in *NPM1wt* and *NPM1c+* AML cells

As we cultured the cells, we noted that we could not maintain the *NPM1c+* caspase-2-deficient cells in culture for longer than three weeks. This was surprising because caspase-2-deficient cells are known to proliferate faster (Boice et al., 2022). To determine if this represented a novel consequence of NPM1c+-dependent caspase-2 activation, we investigated the impact of the loss of caspase-2 on cell cycle progression in *NPM1wt* and *NPM1c*+ cells. Using BrdU/7-AAD staining and flow cytometry, we profiled each cell type to quantitate the proportion of cells in each phase of the cell cycle. Parental *NPM1wt* cells exhibited normal cell cycle progression with the expected horseshoe pattern of BrdU/7-AAD staining showing the three distinct cell cycle phases, G1, S, and G2 (Figure 3A). Loss of caspase-2 in *NPM1wt* cells had a minimal impact on the cell cycle profile compared to the parental cells (Figure 3A, B). In stark contrast, caspase-2 loss in *NPM1c+* cells resulted in a profound increase in cells in G1-phase (Figure 3A, B). These data suggest that caspase-2 is required for normal cell cycle progression of *NPM1c+* AML cells. To investigate this further, we measured cell proliferation over three weeks in freshly thawed cells using an MTT assay. While *NPM1wt* parental, *NPM1wt* caspase-2-deficient, and *NPM1c+* parental cells proliferated as expected (Figure 3C, D), caspase-2-deficient *NPM1c+* cells showed no growth and a decline in viability (Figure 3D). This result provides strong evidence that caspase-2 is required for the sustained proliferation of *NPM1c+* cells.

**Figure 3:**
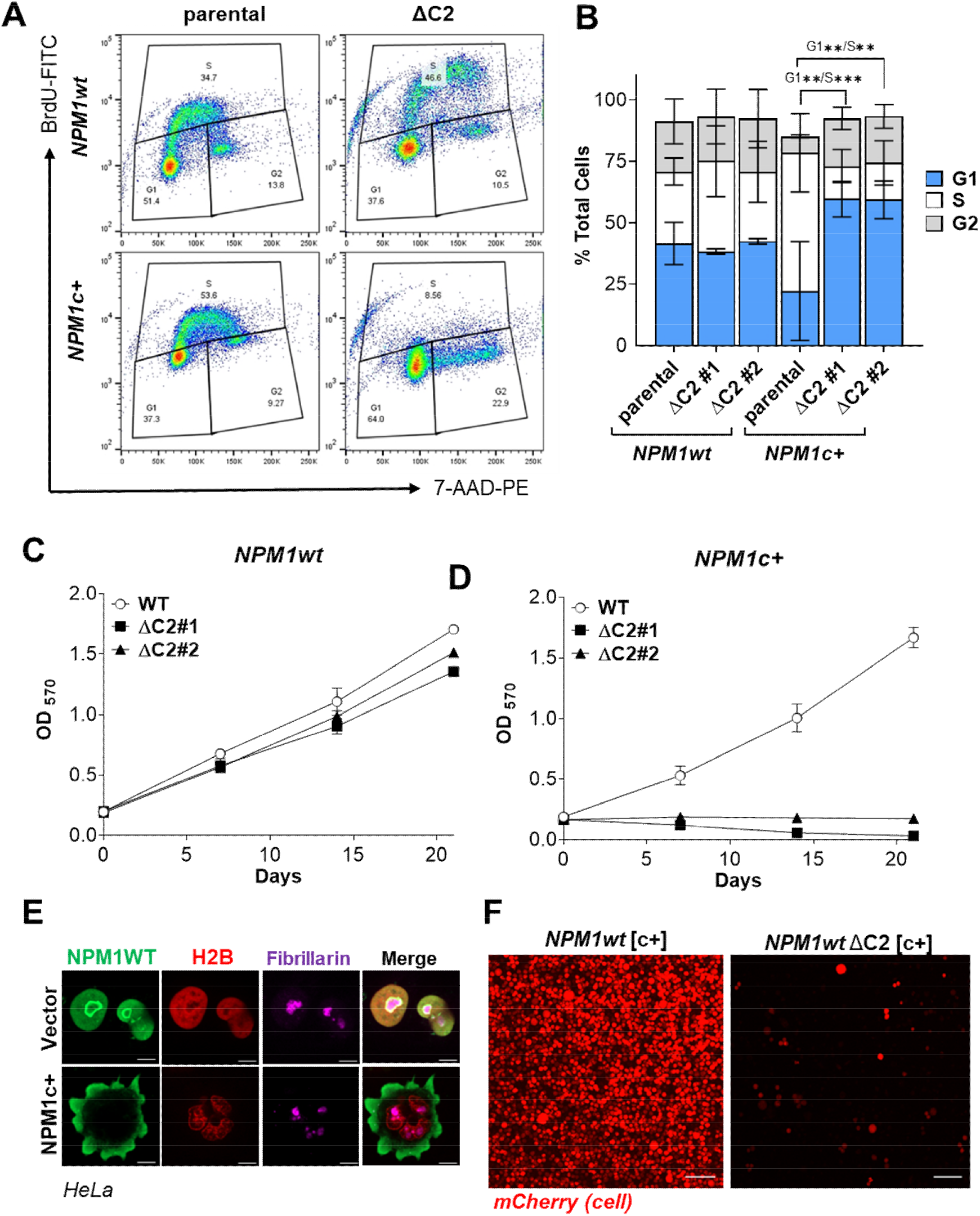
**Loss of caspase-2 induces growth arrest in NPM1c+ cells.** (A) OCI-AML-2 (*NPM1wt*) and OCI-AML-2 (*NPM1c+*) parental or ΔC2 cells were harvested following a 30 min BrdU (10 µM) pulse. The proportion of cells in S-, G1-, and G2-phase was determined by flow cytometry. Representative flow plots are shown. (B) The percent of cells in each phase of the cell cycle was determined for each cell line. Results are the average of two independent experiments plus or minus standard deviation **p < 0.01, ***p < 0.001. (C–D) Cells of each genotype were seeded at 1 x 10^4^ cells/well and cell viability was measured by MTT at the indicated times for *NPM1wt* parental and ΔC2 cells (C) and *NPM1c+* parental and ΔC2 cells (D). Results are the average of three independent experiments plus or minus standard deviation. (E) HeLa cells were transiently transfected with an empty vector or an expression construct for NPM1c+ along with H2B mCherry as transfection reporter. Cells were stained with an anti-fibrillarin antibody to show the nucleolus (*purple*) and anti-B23 antibody (*green*) that recognizes only wild-type NPM1 (see Supplemental Figure S3A). H2B-mCherry fluorescence delineates the nucleus (*red*), Representative images taken 24 h post-transfection are shown. Bar, 10 µm. (F) *NPM1wt* and *NPM1wt*ΔC2 cells transduced with NPM1c+ linked to an mCherry reporter ([c+]) were imaged 2 weeks after antibiotic selection. The viable cells are shown in red. Bar, 10 µm.

To test whether the loss of viability was specific to cells harboring the *NPM1c+* mutation, we reconstituted NPM1c+ expression and function in *NPM1wt* cells. *NPM1c+* is always a heterozygous mutation and, upon expression, it hetero-oligomerizes with the wild-type copy of NPM1 and translocates it from the nucleolus to the cytoplasm (Brodska et al., 2017). Hence, we expected that exogenously expressed *NPM1c+* would export endogenous NPM1 to the cytoplasm, functionally mimicking *NPM1c+* cells. To test this hypothesis, we transiently expressed wNPM1 with a TCTG insertion mutation in exon 12 (NPM1c+) in HeLa cells that normally express wild-type NPM1. We immunostained the cells with a C-terminal NPM1 antibody that only detects wild-type NPM1 and not the mutant form (see Supplemental Figure S3B). As expected, in control cells, wild-type NPM1 was detected in the nucleolus, forming a ring around the nucleolar protein fibrillarin, and in the nucleus, co-localizing with the transfected nuclear marker H2B mCherry (Figure 3E). In cells transfected with NPM1c+, we detected wild-type NPM1 in the cytoplasm of cells. The NPM1 staining was non-nuclear and overlapped with phalloidin, which stains actin as a marker of the cytoplasm (Figure 3E, Supplemental Figure S2). This finding indicates that NPM1c+ induces cytoplasmic translocation of wild-type NPM1.

To efficiently convert *NPM1wt* AML cells to *NPM1c+* cells, we expressed NPM1c+ linked to an mCherry reporter via lentiviral transduction in *NPM1wt* cells. Parental *NPM1wt* cells expressing NPM1c+ (*NPM1wt*[c+]) showed normal proliferation. However, when we expressed NPM1c+ in *NPM1wt* caspase-2-deficient cells (*NPM1wt*[c+] ΔC2), we noted a rapid loss of viability and we could not establish a stable line (Figure 3F). These results demonstrate that the loss of viability specifically in cells lacking caspase-2 is due to the presence of NPM1c+ and not any other genetic difference between the *NPM1wt* and *NPM1c+* cell lines. This result confirms that *NPM1c+* cells require caspase-2 for growth and proliferation.

### Loss of caspase-2 induces NPM1c+-dependent terminal differentiation

Upon depletion from the cytoplasm of AML cells, *NPM1c+* has been shown to induce cell cycle arrest in G1 and terminal differentiation (Brunetti et al., 2018). Given that deletion of caspase-2 induces extended G1 arrest, specifically in *NPM1c+* cells (Figure 3B), we investigated if these cells undergo differentiation in the absence of caspase-2. We measured the expression of the cell surface markers CD34, to measure immature hematopoietic precursors, and CD14, to measure myeloid differentiation. CD34 expression was minimal in the NPM1wt and NPM1c+ cells, as previously reported (Hayun et al., 2020; Pianigiani et al., 2022), and was not changed in the absence of caspase-2 (Supplemental Figure S3A). In contrast, CD14 surface expression was increased more than 10-fold in caspase-2-deficient *NPM1c+* cells (Figure 4A, B). These data indicate that loss of caspase-2 induces differentiation of *NPM1c+* cells into a macrophage-like state and that these cells can no longer proliferate.

**Figure 4:**
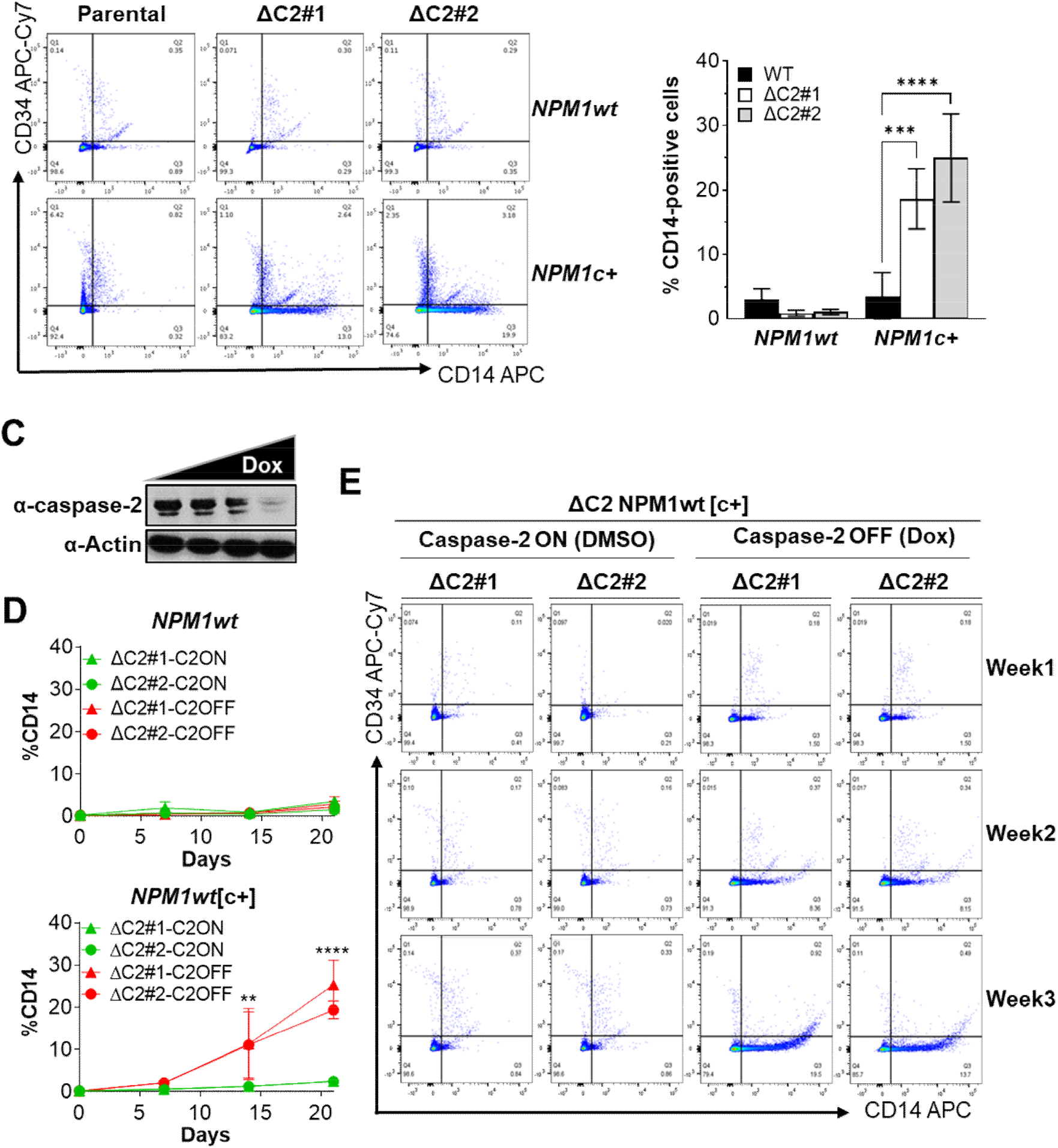
**Loss of caspase-2 induces NPM1c+-dependent differentiation.** (A) Cell surface expression of CD34 and CD14 was measured in OCI-AML-2 (*NPM1wt*) and OCI-AML-3 (*NPM1c+*) parental or ΔC2 cells cultured for two weeks by flow cytometry. Representative flow plots are shown. (B) The percentage of cells expressing CD14 is shown. Results are the average of three independent experiments plus or minus standard deviation, ***p < 0.001, ****p < 0.0001. (C) *NPM1wt*ΔC2[c+] cells expressing a Tet-repressible caspase-2 (TetOFF-caspase-2) were treated with increasing doses of doxycycline (0.1–2 µg/mL) for 16 h to repress caspase-2 expression. Lysates were immunoblotted for caspase-2 and actin as a loading control. (D) *NPM1wt*[c+]-C2^OFF^cells were grown in DMSO (C2ON) or doxycycline (2 µg/mL, C2OFF) for 3 weeks. Cell surface CD14 and CD34 expression was measured once a week by flow cytometry. Results represent the average of three independent experiments, **p < 0.01, ****p < 0.0001. (E) Representative flow plots from (D) are shown (see also Supplemental Figure S3C).

To demonstrate that caspase-2 is required for this process and no other genetic difference between the cell lines, we reintroduced caspase-2. We used a TET-OFF-based expression system where we could both reconstitute caspase-2 expression and repress caspase-2 transcription in the same cells. Due to the reduced viability of caspase-2-deficient *NPM1c+* cells, they did not survive long enough to establish a stable cell line. To circumvent this, we stably expressed the repressible caspase-2 in *NPM1wt* caspase-2-deficient cells (*NPM1wt* ΔC2) and then converted them to *NPM1c+* by expressing exogenous *NPM1c+* as shown in Figure 3F. We will refer to these cells as *NPM1wt*[c+]C2^TET^. Unlike the *NPM1wt*[c+]ΔC2 cells (Figure 3F), the *NPM1wt*[c+]C2^TET^ survived selection and had no obvious growth impairment. This indicates that the caspase-2 re-expression prevents the loss of viability associated with expression of NPM1c+ in caspase-2-deficient cells. The addition of doxycycline to the *NPM1wt*[c+]C2^TET^ cells efficiently inhibited caspase-2 expression in a dose-dependent manner (Figure 4C, Supplemental Figure S3B). We assessed the repressible caspase-2 cells for CD34 and CD14 surface expression under caspase-2 ON (grown in the absence of doxycycline) and caspase-2 OFF (grown in the presence of doxycycline) conditions over a three-week period. As expected, neither restoration of caspase-2 (C2ON) nor depletion of caspase-2 after doxycycline treatment (C2OFF) affected the percentage of CD14-positive cells in the *NPM1wt* lines (Figure 4D, Supplemental Figure S3C).

In *NPM1wt*[c+] cells with cytoplasmic expression of NPM1, *NPM1wt*[c+]C2^TET^ cells grown in DMSO (C2ON) did not differentiate to express CD14 over the three weeks (Figure 4D, E). This confirms that the reintroduction of caspase-2 to the cells restores them to the *NPM1c+* parental phenotype. Depletion of caspase-2 with doxycycline (C2OFF) in the *NPM1wt*[c+]C2^TET^ cells resulted in a time-dependent increase in CD14 surface expression, peaking at week 3 (Figure 4D, E). Taken together, these results confirm that caspase-2 is required to sustain viability by preventing the monocytic differentiation of *NPM1c+* cells.

### Caspase-2 loss downregulates genes regulating stem cell pluripotency

To further understand the molecular mechanisms by which caspase-2 sustains *NPM1c+* cell proliferation and prevents differentiation, we performed transcriptomic analysis. We carried out total RNAseq on parental *NPM1wt*, *NPM1c+* cells, and their respective caspase-2-deficient counterparts (Figure 5A). By comparing the differentially expressed genes with greater than 5- fold expression changes across the genotypes, we noted that caspase-2 deficiency had an extensive impact on the transcriptomic profile of *NPM1c+* cells but not *NPM1wt* cells (Figure 5B). Given this large difference in expression between *NPM1c+* cells with and without caspase-2, we focused on the differences between these cell lines. We used gene enrichment analysis to first determine cell types associated with the changes in gene expression after loss of caspase-2 in *NPM1c+* cells. We found that the changes in expression profile were associated with an increased identity with CD14+ monocytes according to the Human Gene Atlas (Supplemental Figure S4A). There were significant increases in the macrophage markers CD14, CD86, and CYBB (Supplemental Figure S4B). This confirms our observations that loss of caspase-2 induces monocytic differentiation of *NPM1c*+ cells. Next, we carried out KEGG pathway analysis. We found that the most enriched pathways between *NPM1c*+ parental and *NPM1c*+ΔC2 cells were “Pathways in Cancer,” “Hippo Signaling Pathway,” and “Signaling Pathways Regulating Pluripotency of Stem Cells” (Figure 5C). In fact, the most significantly downregulated group was Signaling Pathways Regulating Pluripotency of Stem Cells (Table 1). In particular, the FGF2 pathway (*FGF2*, *FGFR*, *AKT3*), the IGFR pathway (*IGF1R*, *AKT3*), the Wnt pathway (*WNT3*, *WNT5B*, *WNT7B*, *WNT11*, *FZD1–4*, *TCF7*), the LIF pathway (*LIF*, *LIFR*, *JAK3*) and one of the core members of the self-renewal transcriptional network, Oct4 (*POU5F1B*) were all downregulated in the absence of caspase-2 in *NPM1c+* cells (Figure 5D). Together, these data provide strong evidence that caspase-2 is required to maintain *NPM1c+* AML self-renewal.

**Figure 5:**
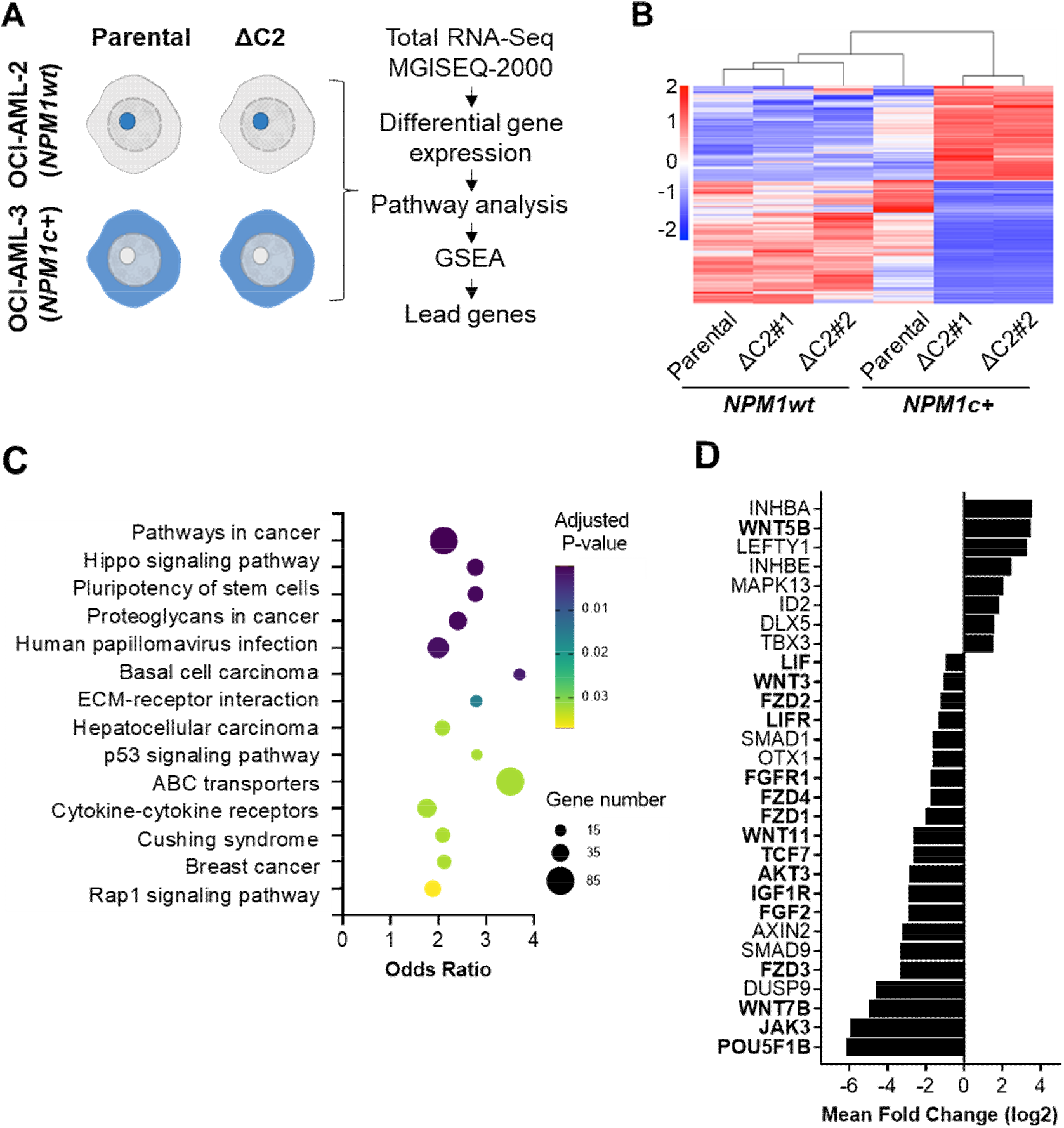
**Loss of caspase-2 in *NPM1c+* cells downregulates pathways regulating pluripotency.** (A) The experimental design for total RNA sequencing and analysis of differentially expressed genes (DEGs) from parental OCI-AML-2 (*NPM1wt*) and OCI-AML-3 (*NPM1c+*) and their respective caspase-2-deficient (ΔC2) cells is shown. (B) DEGs between the parental *NPM1wt* and *NPM1c+* and their respective ΔC2 transcriptomes are shown by heat map. Genes with more than log5-fold changes are shown. Decreased expression is shown in blue and increased expression is shown in red. (C) Total RNAseq data were analyzed with ENRICHR to identify genes involved in the KEGG term biological processes indicated that are most significantly differentially regulated in *NPM1c+* parental and ΔC2 cells. The size of the dots represents the number of genes associated with each biological pathway. The color of the dots represents the significance (adjusted p-value). (D) The mean log2 fold change of significant DEGs between *NPM1c+* parental and ΔC2 cells involved in the KEGG term pathways that regulate pluripotency of stem cells (the most significantly downregulated pathway (see Table 1)) is shown. Gene names in bold are referred to in the text.

**Table 1:** Most significantly up and downregulated pathways in NPM1c+ cells in the absence of caspase-2.

|  | Term | Overlap | P-value | Adjusted P-value | Odds Ratio | Combined Score |
| --- | --- | --- | --- | --- | --- | --- |
| Down regulated | Signaling pathways regulating pluripotency of stem cells | 21/143 | 5.93E-06 | 0.001671943 | 3.414446184 | 41.09516759 |
|  | Hippo signaling pathway | 21/163 | 4.5E-05 | 0.006338397 | 2.930434654 | 29.33332658 |
|  | Pathways in cancer | 46/531 | 0.000117 | 0.008361907 | 1.894965339 | 17.15021715 |
|  | Human papillomavirus infection | 32/331 | 0.000183 | 0.008361907 | 2.127629963 | 18.31472613 |
|  | Cushing syndrome | 19/155 | 0.000198 | 0.008361907 | 2.763396982 | 23.5675838 |
|  | Hepatocellular carcinoma | 20/168 | 0.000203 | 0.008361907 | 2.674089184 | 22.74221776 |
| Upregulated | Malaria | 10/56 | 9.92624E-06 | 0.002729716 | 6.745091164 | 77.70566909 |
|  | Pathways in cancer | 39/531 | 2.41339E-05 | 0.003318406 | 2.176980816 | 23.14543067 |
|  | Cytokine-cytokine receptor interaction | 25/295 | 7.58836E-05 | 0.006956 | 2.521357317 | 23.91837579 |
|  | Viral protein interaction with cytokine and cytokine receptor | 12/100 | 0.000259819 | 0.017862524 | 3.680283851 | 30.38268317 |
|  | Toll-like receptor signaling pathway | 12/104 | 0.000375033 | 0.019012637 | 3.5195377 | 27.76386473 |

We next confirmed that these changes in gene expression resulted in protein changes. We did so in an unbiased manner by using reverse phase protein array (RPPA) analysis of the *NPM1c+* parental and caspase-2-deficient cells. Of the protein set tested, we found that the majority of proteins with significant differences between the two cell lines (q < 0.01) had decreased abundance in the absence of caspase-2 (Figure 6A). Many of these tested proteins that had decreased abundance in *NPM1c+* caspase-2-deficient cells are known to be required for or support hematopoiesis (Figure 6A). These include a number of epigenetic modifiers like JMJD2A, JMJD2B, and LSD1, that have all been shown to be required for stem cell maintenance (Agger et al., 2019; Hu et al., 2009) (Table 2). Of the proteins that were significantly changed, only one, HIF2A, was increased in the absence of caspase-2. In the largest group, fifteen of the most significantly down-regulated proteins are involved in the AKT/mTORC1 pathway, and three are involved in the Wnt pathway, including β-catenin, the downstream effector of Wnt signaling (Table 2). These findings are consistent with the RNAseq data that show downregulation of pathways that converge on AKT and Wnt signaling (Figure 5D).

**Figure 6:**
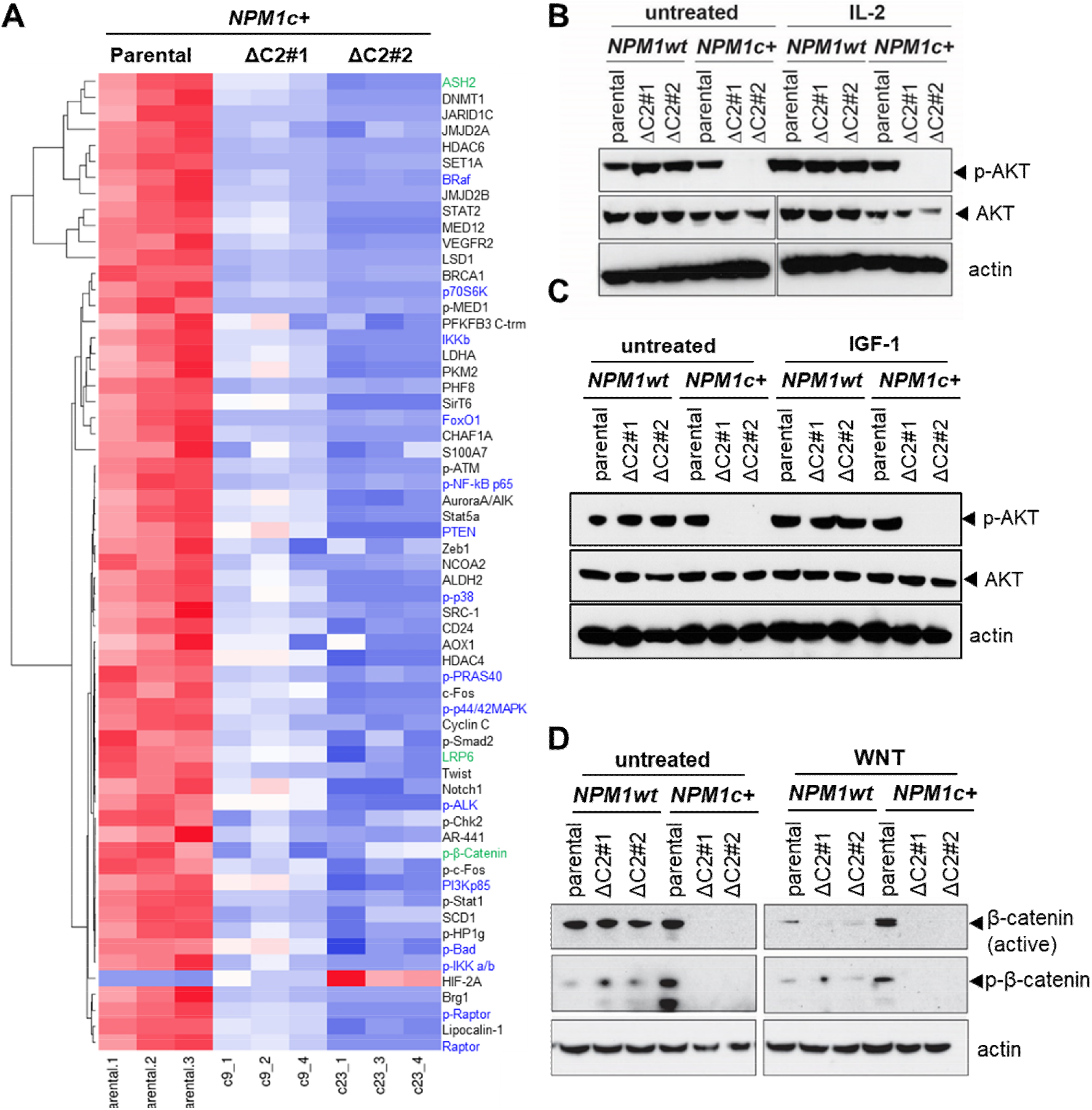
**Caspase-2 regulates pathways that maintain stemness in *NPM1c+* cells.** (A) Reverse phase protein array (RPPA) analysis was carried out on OCI-AML2 (*NPM1c+*) parental and ΔC2 cells. Signal intensities were normalized and filtered as described in Materials and Methods. The heat map shows Z-scores of three biological replicates of parental *NPM1c+* cells and each ΔC2 clone with q > 0.01. Upregulated proteins are shown in red and down regulated proteins are shown in blue. Proteins in the mTORC1/AKT pathway are labeled in blue and proteins in the Wnt signaling pathway are labeled in green. (B) OCI-AML-2 (*NPM1*wt) and OCI-AML-3 (*NPM1c*+) parental or ΔC2 cells were left untreated or treated with IL-2 for 16 h. Lysates were harvested and immunoblotted for phosphorylated AKT (p-AKT) and total AKT. Actin was used as the loading control. A spacer lane was cropped from the actin and AKT blots for alignment purposes. (C) *NPM1wt* and *NPM1*c+ parental or ΔC2 cells were treated with or without IGF-1 (10 ng/mL) for 16 h. Lysates were harvested and immunoblotted for p-AKT and total AKT. Actin was used as the loading control. (D) *NPM1wt* and *NPM1*c+ parental or ΔC2 cells were treated with or without Wnt (5.0 nM) for 16 h. Lysates were harvested and immunoblotted for β- catenin (active) and phosphorylated β-catenin (inactive). Actin was used as the loading control. The panels for each protein are from the same immunoblot with irrelevant lanes cropped out. Experiments shown in (B–D) are representative of 2–3 independent experiments.

**Table 2:**
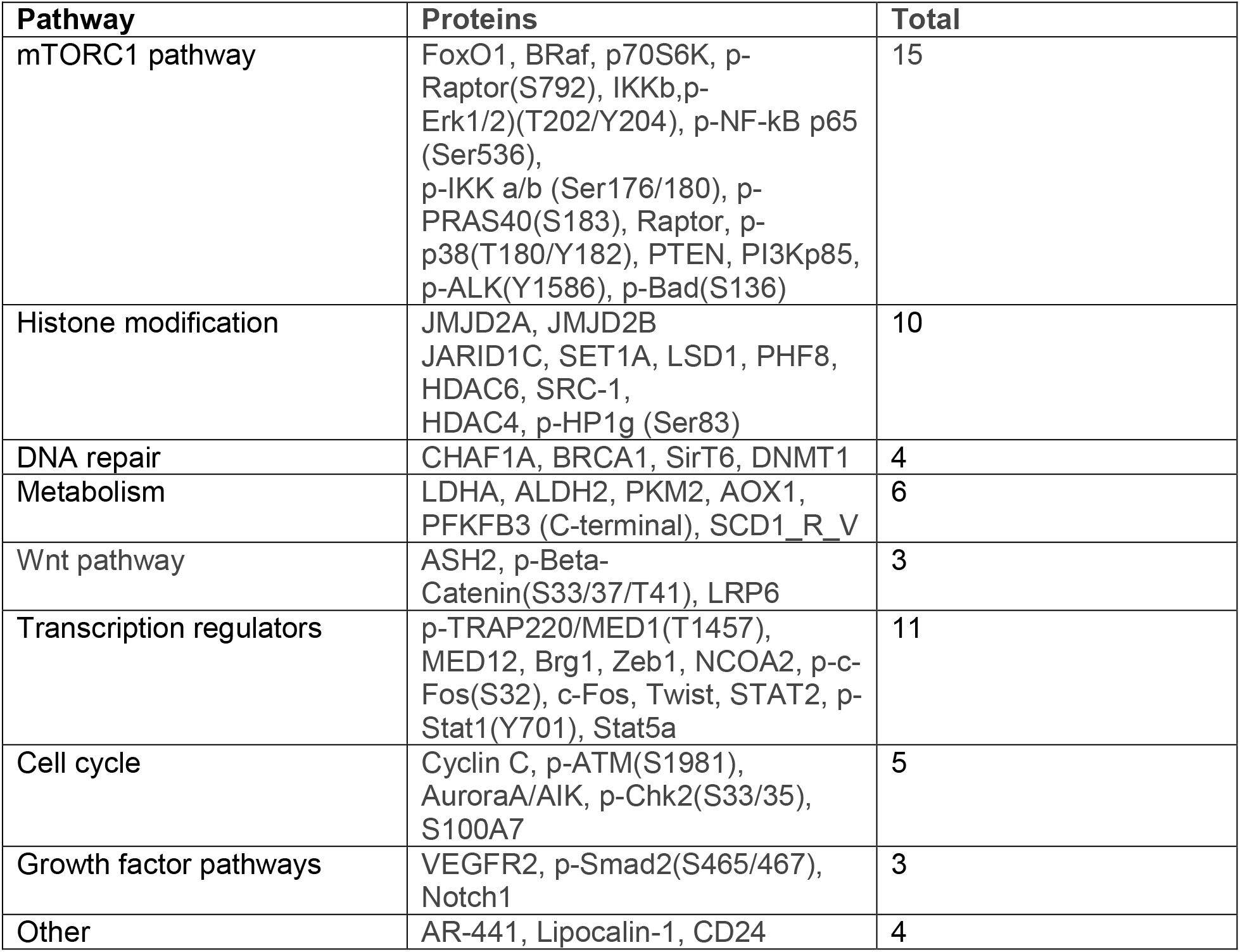
Significantly downregulated proteins in *NPM1c+* cells in the absence of caspase-2.

The FGFR, IGF1R, and LIFR pathways were all downregulated in the RNAseq data set in the absence of caspase-2. These pathways all converge on activation of the PI3K pathway resulting in phosphorylation of AKT. Because the AKT pathway featured heavily in the RPPA results, we tested the impact of the loss of caspase-2 on AKT activation. Unfortunately, p-AKT was excluded from the RPPA data set based on the filtering criteria. Instead, we measured the effects of the loss of caspase-2 on general AKT activation using IL-2 as a stimulus. IL-2 induced phosphorylation of AKT in both the *NPM1wt* and *NPM1c+* parental cells. Loss of caspase-2 had minimal effect on the amount of AKT phosphorylation in *NPM1wt* cells. In contrast, AKT phosphorylation was completely inhibited in the *NPM1c+* caspase-2-deficient cells (Figure 6B). Next, we treated the cells with human IGF-1 recombinant protein. Following IGF-1 treatment, we observed phosphorylation of AKT in the *NPM1wt* and *NPM1c*+ parental lines but none in the caspase-2-deficient *NPM1c+* cells after IGF-1 treatment (Figure 6C). Total AKT levels were similar in all cell types. Therefore, the lower phosphorylation of AKT cannot be explained just by lower AKT levels. We also noted an increase in the basal level of AKT phosphorylation in caspase-2-deficient NPM1wt cells (Figure 6B, C). This suggests that caspase-2 has opposite effects on the *NPM1wt* cells compared to the *NPM1c+* cells. These data confirm that caspase-2 loss impairs the AKT pathway, specifically in *NPM1c+* cells.

Another downregulated pathway that was common between the RNAseq and RPPA data was the Wnt signaling pathway. Therefore, we confirmed the effect of loss of caspase-2 on Wnt signaling. The downstream effector of the canonical Wnt pathway is β-catenin, the constant degradation of which is induced when it is phosphorylated. Wnt induces enhanced transcriptional activation of and stabilization of β-catenin. (Clevers and Nusse, 2012). We treated the AML cells with the Wnt surrogate-Fc fusion recombinant protein, which stabilizes β-catenin. In untreated cells, we detected both phosphorylated (inactive) and unphosphorylated (active) β-catenin in *NPM1wt* cells with and without caspase-2 and in *NPM1c*+ cells (Figure 6D). Phosphorylated β-catenin was increased in the parental *NPM1c*+ cells but both forms of β-catenin were absent in the *NPM1c+*ΔC2 cells. Following treatment with Wnt, overall levels of phospho-β-catenin were reduced in all *NPM1wt* cell types and in *NPM1c+* parental cells compared to the untreated cells. In contrast, the active form of β-catenin was high in *NPM1c+* parental cells. In the absence of caspase-2, β-catenin and phosphorylated β-catenin were both absent. This indicates that the expression or stabilization of β-catenin is impaired in the absence of caspase-2 in *NPM1c+* AML. This is consistent with the reduced expression of the upstream frizzled receptors in the Wnt signaling pathway in *NPM1c+* caspase-2-deficient AML cells (Figure 5D).

## Discussion

*NPM1c+* is the most frequent genetic alteration in AML and, as such, represents a potential therapeutic target (Falini et al., 2005). Despite this importance, how this driver mutation induces and maintains the leukemic state is not clear. The data presented here demonstrate that the cellular effects and signaling mechanisms of *NPM1c+* converge on the pro-apoptotic protein caspase-2. We show that caspase-2 is a crucial link between NPM1c+ and changes in oncogene signature, without which the cells undergo significant differentiation and lose their proliferative and self-renewing capacity. Moreover, our results show that caspase-2 activation has vastly different functional outcomes depending on the specific upstream signals.

Building on our prior discovery that NPM1 is an upstream activator of caspase-2 in the nucleolus (Ando et al., 2017), we show here that NPM1c+ has a similar ability to activate caspase-2 in the cytoplasm. Surprisingly, this leads to vastly different functional consequences than would be expected. Instead of enhancing proliferation as has been previously reported (Boice et al., 2022; Ho et al., 2009), loss of caspase-2 in *NPM1c*+ cells induced profound cell cycle arrest and what appears to be terminal differentiation. These findings provide strong evidence that caspase-2 is essential for survival of *NPM1c*+ AML cells. The dramatic impact on the stem cell transcriptional signature converging mainly on the AKT/mTORC1 and Wnt signaling pathways indicate the essential role for caspase-2 in survival is not solely due to a direct regulation of the cell cycle but rather that caspase-2 plays an integral role in maintaining the self-renewing capacity of *NPM1c+* AML cells.

The few studies that have investigated the role of caspase-2 in hematopoiesis have shown that caspase-2 serves to limit this process. For example, in aged mice, the absence of caspase-2 increased the number of myeloid progenitor cells and impaired the differentiation of hematopoietic stem cells (Dawar et al., 2016). In addition, hematopoietic cell lineage pathways were found to be significantly upregulated in *Casp2*^-/-^ Eµ-Myc lymphoma compared to *Casp2*^+/+^ Eµ-Myc (Dorstyn et al., 2019). When NPM1c+ is expressed, however, we found that differentiation was increased, and stem cell pathways were downregulated in the absence of caspase-2. These results suggest that, instead of leading to inhibition of hematopoiesis, caspase-2 activation by NPM1 outside of the nucleolus leads to the promotion of pluripotency. This opposite action of NPM1-induced caspase-2 activation in the cytoplasm versus the nucleolus is supported by the small changes we see in the *NPM1wt* cells. For example, there is a small increase in the amount of DNA damage-induced caspase-3 cleavage in NPM1wt caspase-2-deficient cells, even though this does not result in a short-term impact on apoptosis. We also observed an increase in basal AKT activation in *NPM1wt* caspase-2-deficient cells. Our results support a model where caspase-2 has opposing functions depending on where it is activated in the cell. Since caspase-2 is a protease, it is likely that the compartmentalization of caspase-2 activation into different regions of the cell provides access to different substrates.

The functional compartmentalization of caspase-2 due to differential substrate access is demonstrated by the canonical function of caspase-2 in inducing apoptosis in *NPM1c+* AML cells. Following DNA damage, we only saw caspase-2-dependent apoptosis in the *NPM1c+* cells. Caspase-2 induces apoptosis by inducing BID cleavage (Bonzon et al., 2006; Upton et al., 2008). The improved ability for caspase-2 to be able to induce apoptosis in *NPM1c+* cells is likely due to its closer proximity to BID in the cytoplasm once it is activated. Interestingly, apoptosis is not completely inhibited in the absence of caspase-2 in *NPM1c+* cells but is rather reduced to the same level as that observed in the *NPM1wt* cells. This result indicates that caspase-2 is responsible for the increased sensitivity of these cells, but other apoptotic pathways also contribute to the cell death.

How caspase-2 both induces apoptosis and proliferation in the same cell type appears to be somewhat of a conundrum. The key element here appears to be the stimulus. The caspase-2 knockout *NPM1c+* cells show short-term resistance to apoptosis when challenged with a DNA damaging agent, but in the absence of an external stimulus, will arrest, differentiate, and eventually lose viability. It is possible that the enhanced self-renewal capacity and rapid proliferation of the NPM1c+ cells (as demonstrated by increased S-phase cells) make the cells more sensitive to pro-apoptotic stimuli. Indeed, pluripotent stem cells have been shown to have increased sensitivity to apoptosis induced by genotoxic stimuli (Smith et al., 2012). This mechanism been proposed to be a way to limit the accumulation of DNA damage and genomic instability (Liu et al., 2014).

It has been proposed that NPM1c+-mediated expression of HOX genes is required for the maintenance of the leukemic state of the AML cells (Brunetti et al., 2018). Interestingly, we did not observe large differences in HOX gene expression in *NPM1c+* cells with or without caspase-2. The C-terminus of NPM1 is comprised of a cluster of helices (Helix 1–3) that bind both duplex and single-stranded DNA with no sequence-specificity (Gallo et al., 2012). When mutated, Helix 1 becomes denatured and unstructured, which impacts the ability of NPM1 to bind DNA and protein (Gallo et al., 2012). NPM1c+ is recruited to the HOX gene cluster by CRM1 via binding to the newly formed NES in its sequence (Oka et al., 2019). Disruption of the NES or depletion of NPM1c+ from the cytoplasm disrupts this interaction (Brunetti et al., 2018). Because NPM1c+ is thought to directly impact HOX expression, it is perhaps not surprising that they are not disrupted in caspase-2-deficient cells. Our data suggest that caspase-2 is another downstream effector of NPM1c+ that helps maintain pluripotency. The role of AKT in modulating the expression of core stem cell regulators, OCT4, SOX2, and Nanog is described in multiple studies (Malak et al., 2015; Schaefer et al., 2015; Yoon et al., 2021). Such studies advocate that activation of AKT signaling is sufficient to maintain stem cell pluripotency (Watanabe et al., 2006). In addition, constitutive AKT activation transforms hematopoietic stem cells into myeloproliferative diseases, including AML (Kharas et al., 2010). AML cells often have hyper-activated AKT, a prerequisite to sustain the oncogenic potential of leukemia stem cells and prevent differentiation (Bertacchini et al., 2014; Park et al., 2010). Consistent with our data, NPM1wt and NPM1c+ have opposing effects on AKT signaling, as it has been shown that NPM1 can negatively regulate AKT and NPM1c+ antagonizes NPM1-mediated suppression of AKT signaling (Ren et al., 2020). The relationship between AKT and HOX function is not clear, but it has been suggested that the HOXD gene cluster induces AKT activation (Wang et al., 2022a). Therefore, it is possible that caspase-2 functions in synergy with HOX genes to engage the AKT/mTORC1 pathway.

It is unlikely that AKT is the only effector of NPM1c+-mediated caspase-2 activation. Caspase-2 also engages the WNT-signaling pathway in *NPM1c+* cells, as evidenced by reduced expression of the Frizzled receptors and reduced β-catenin expression and activation. Among the differentially expressed Wnt-signaling genes between NPM1c+ parental and caspase-2-deficient cells, WNT5b is upregulated in the absence of caspase-2. This ligand is antagonistic to the canonical WNT signaling pathway (Suthon et al., 2021), fitting with the overall downregulation of this pathway in the absence of caspase-2. Because a number of receptors in the stem cell pathways upstream of AKT and β-catenin were downregulated in the absence of caspase-2, we predict that caspase-2 functions upstream of these receptors, likely by cleaving a substrate(s) that regulate their transcription. It has been notoriously difficult to identify bona fide caspase-2 substrates (Brown-Suedel and Bouchier-Hayes, 2020). Although a large-scale caspase substrate screen has been published identifying a long list of potential caspase-2 substrates (Julien et al., 2016), the only two that have been shown to have clear physiological consequences when cleaved by caspase-2 are BID and the p53 negative regulator MDM2 (Oliver et al., 2011). More work is needed to determine the specific targets of caspase-2 in the novel pathway described here.

The high rate of relapse, minimal residual disease, and associated mortalities in AML underscores the need to develop targeted therapies. In the absence of additional mutations, *NPM1c+* AML is associated with high rates of complete molecular remission following standard induction therapy (Falini et al., 2007). However, when combined with one or more mutations like *FLT3-ITD*, *DNMT3a*, or *IDH1*, prognosis is worse (Dovey et al., 2017), and overall relapse-free survival is the same as in patients with *NPM1wt* AML (Issa et al., 2023). The self-renewal properties of AML cells provide a potential reservoir for relapse. Despite our studies suggesting that the AKT pathway is a major convergence point in *NPM1c+* AML pluripotency, clinical studies with AKT pathway inhibitors as monotherapy for *NPM1c+* AML showed insufficient anti-leukemic activity (Konopleva et al., 2014). The poor efficacy of AKT inhibitors for leukemia treatment underscores the importance of understanding the full extent of the pathways engaged by NPM1c+ and caspase-2. Our work here holds the promise that targeting caspase-2 in *NPM1c+* AML could be an effective method to treat the disease and prevent relapse.

## Materials and Methods

### Chemicals and Antibodies

The following antibodies were used: anti-NPM1 (clone FC82291; sigma-aldrich); anti-caspase-2 (clone 11B4; EMD Millipore); anti-fibrillarin (C13C3; Cell Signaling Technology); anti-BID (AF860; R&D systems); anti-caspase-3 (9662; Cell Signaling Technology); anti-caspase-3 cleaved (9661; Cell Signaling Technology); anti-Actin (clone 4; Fisher Scientific); anti-phospho-AKT (4060; Cell Signaling Technology), anti-AKT (4691; Cell Signaling Technology); anti-β-catenin (active) (8814; Cell Signaling Technology); anti-β-catenin (inactive) (961; Cell Signaling Technology); anti-B23 (172-6195; Fisher-scientific); APC/Cy7 anti-human CD34 (clone 581; BioLegend); APC anti-human CD14 (clone 63D3; BioLegend). Alexa Fluor 647 Phalloidin was purchased from Cell Signaling Technology. All cell culture media reagents were purchased from Thermo Fisher Scientific. Unless otherwise indicated, all other reagents were purchased from Sigma-Aldrich (St. Louis, MO, USA).

### Plasmids

The caspase-2 BiFC reporter pRRL-C2 Pro-VC-2A-C2 Pro-VN 2A-mCherry-2A-Puro was described previously (Ando et al., 2017). The pcDNA3.1.NPM1c+ plasmid was generated by site-directed mutagenesis of pcDNA3.1-NPM1 using the primer sequences FWD: 5’- CTATTCAAGATCTCTGTCTGGCAGTGGAGGAAGGCG-3’ and REV: 5’-CTATTCAAGATC<u>TCTG</u>TCTGGCAGTGGAGGAAGGCG-3’ to insert the TCTG nucleotide in exon 12 of the NPM1 gene between sites 1864-1868. NPM1c+ was cloned into pRRL-MND-MCS- 2A-mCherry-2A-Puro by standard PCR strategies with primers designed to incorporate restriction enzymes, XhoI and NotI. pLVX-Tet-Off Advanced and pLVX Tight-Puro-GOI vectors were purchased from Takara Bio (cat#632163) as a part of the Lenti-X Tet-off Advanced Inducible Expression system. Caspase-2 was cloned into the pLVX Tight-Puro vector by the standard PCR strategies. Histone H2B-mCherry was purchased from Addgene (#20972 (Nam and Benezra, 2009)). Each construct was verified by sequencing.

### Cell culture and generation of cell lines

OCI-AML2 (*NPM1* WT) and OCI-AML3 (*NPM1c+*) cells were grown in RPMI containing FBS (10% v/v), L-glutamine (2 mM), and penicillin/streptomycin (50 IU/mL/50 µg/mL). Human Embryonic Kidney (HEK) 293T and HeLa cells were grown in DMEM containing FBS (10% v/v), L-glutamine (2 mM), and penicillin/streptomycin (50 IU/mL/50 µg/mL). Stable cell lines were generated by lentiviral transduction. HEK 293T cells were transiently transfected with pRRL.C2 Pro-VC-2A-C2 Pro-VN-2A-mCherry along with the packaging vector, pSPAX-2, and the envelope vector pVSV-G using Lipofectamine 2000 transfection reagent (Thermo Fisher Scientific) according to manufacturer’s instructions. Additional lentiviral vectors were packaged in HEK 293T cells using Lenti-X Packaging Single Shots (VSV-G) (Takara Bio USA) per the manufacturer’s protocol. After 48 h, virus-containing supernatants were cleared by centrifugation and incubated with OCI-AML2 or OCI-AML3 cell lines with polybrene (5 µg/mL). For the Tet-repressible cell lines, OCI-AML cell lines were co-transduced with pLVX-Tet-Off Advanced and pLVX Tight-Puro-caspase-2 vectors in a 1:1 ratio. Cells were selected in puromycin (1-5 µg/mL) or G418 (400 µg/mL), followed by bulk fluorescence-activated cell sorting (FACS AriaII, BD) when a linked fluorescent gene was expressed.

### CRISPR/Cas9 gene editing

Caspase-2 was deleted from OCI-AML-2 and OCI-AML-3 cells using an adaptation of the protocol described in (Gundry et al., 2016). Protospacer sequences for caspase-2 were identified using the CRISPRscan scoring algorithm (www.crisprscan.org (Moreno-Mateos et al., 2015)). DNA templates for single-guide RNA (sgRNAs) were made by PCR using a pX459 plasmid containing the sgRNA scaffold sequence and the following primers: ΔCASP2(76) sequence: ttaatacgactcactataGGCGTGGGCAGTCTCATCTTgttttagagctagaaatagc; ΔCASP2(73) sequence: ttaatacgactcactataGGTGTGGAGGGCGCCATCTAgttttagagctagaaatagc; universal reverse primer: AGCACCGACTCGGTGCCACT. sgRNAs were generated by *in vitro* transcription using the HiScribe T7 high-yield RNA synthesis kit (New England Biolabs). Purified sgRNA (0.5 µg) was incubated with Cas9 protein (1 µg, PNA Bio) for 10 min at room temperature. OCI-AML-2 and OCI-AML-3 cells were electroporated with sgRNA/Cas9 complex using the Neon transfection system (Thermo Fisher Scientific) under the condition of 1350 V, 35 ms pulse-width, and one pulse. Single-cell clones were isolated and knockout was confirmed by PCR and western blotting. The single-cell clones were validated by sequencing.

### Flow cytometry

All flow experiments were performed using an LSR Fortessa Flow Cytometer (BD, San Jose, CA, USA). Data were analyzed using FlowJo Software (BD). For Annexin V binding, cells were treated as indicated in the figure legends. Cells were harvested by centrifugation at 300 x g for 5 min and washed with 1X PBS, and resuspended in 200 µL of Annexin V staining buffer (10 mM HEPES, 150 mM NaCl, 5 mM KCl, 1 mM MgCl_2_, and 1.8 mM CaCl_2_) containing 2 µL of Annexin V-APC (Thermo Fisher Scientific). After 15 min of incubation at room temperature, Annexin V-positive cells were quantified by flow cytometry. For cell cycle analysis, cell medium was exchanged for medium with 10 µM BrdU. After 30 min, cells were harvested by centrifugation at 300 x g for 5 min. Cells were fixed and permeabilized with BD Cytofix/Cytoperm buffer for 15 min at room temperature, then with the secondary permeabilization buffer BD Cytoperm Permeabilization Buffer Plus for 10 min on ice, and then were fixed and permeabilized again with BD Cytofix/Cytoperm buffer for 5 min at room temperature. Cells were washed in BD Perm/Wash buffer with FBS and centrifuged at 300 x g between each step. The cells were treated with DNAse (30 µg/1×10^6^ cells) for 1 h at 37°C to uncover the BrdU epitope. Cells were then washed in BD perm/Wash buffer and centrifuged at 300 x g. The cell pellets were incubated in a 1:50 dilution of FITC-labeled anti-BrdU antibody in BD Perm/Wash buffer for 20 min at room temperature. The cells were then resuspended in stain buffer (3% FBS in PBS) containing 10 µL 7-AAD/M cells. BrdU and 7-AAD positive cells were quantitated by flow cytometry. For quantitation of cell surface markers, cells were centrifuged at 1,400 rpm for 5 min. Cell pellets were suspended in 100 µL of fresh media with the appropriate antibodies to cell surface markers and incubated for 45 min at 4°C. The cells were washed in 2 mL of BD Stain Buffer with FBS and centrifuged at 1,400 rpm for 5 min between each step. The cells were fixed and permeabilized with BD Cytofix/Cytoperm buffer for 15 min. Cells were resuspended in 300 µL of stain buffer and quantified for cell surface marker expression by flow cytometry.

### Immunoblotting

Cells were lysed in RIPA buffer (150 mM NaCl, 50 mM Tris-HCL, pH 7, 0.1% SDS, 0.5% sodium deoxycholate, 1% NP-40 (IGEPAL) plus complete protease inhibitors-EDTA (1 mini tablet/10 mL)). 40 µg of cleared protein lysates were resolved by SDS-PAGE on a 4–12% gradient gel (Thermo Fisher Scientific). The proteins were transferred to nitrocellulose membrane (Bio-Rad Laboratories) and immunodetected using relevant primary and peroxidase-conjugated secondary antibodies: Donkey Anti-Rabbit IgG (cat# 84-854, Prometheus), Goat Anti-Rat IgG (cat# 20-307, Prometheus), Sheep Anti-Mouse IgG (cat #84-848, Prometheus), and Donkey Anti-Goat IgG (sc-2020 Santa Cruz). Proteins were visualized with West Dura and West Pico chemiluminescence substrate (Thermo Fisher Scientific).

### Nucleolar isolation

Nucleoli were isolated as per the protocol adapted from (Mitrea et al., 2016). 1 x 10^6^ OCI-AML-2 and OCI-AML-3 cells were washed in 10 mL PBS and centrifuged at 400 x g for 5 min. Cell pellets were resuspended in five pellet volumes of nucleolar isolation buffer (NIB: 10 mM Tris, 2 mM MgCl_2_, 0.5 mM EDTA) containing complete protease inhibitor (Roche). Samples were incubated in NIB buffer for 2 min at room temperature and then on ice for 10 min. Plasma membranes were lysed by the addition of 10% (v/v) IGEPAL to a final concentration of 1% and light vortexing. Crude nuclei pellets were isolated by centrifugation at 500 x g for 3 min. The supernatant was removed and stored as the cytoplasmic fraction. The nuclear pellets were washed in 15 pellet volumes of NIB, 1% IGEPAL. Pellets were then resuspended in 10 pellet volumes of NIB and sonicated at 20% power for 5 cycles of 1 second on followed by 5 seconds off on an Misonix XL 2020 sonicator fitted with a microtip probe (Misonix, Farmingdale, NY, USA). The sonication was repeated until the intact nucleoli were visible. The samples were centrifuged and the supernatant was removed and stored as the nucleoplasmic fraction. The nucleoli pellets were washed once more with 5 pellet volumes of NIB and finally resuspended in 50 µL NIB.

### Microscopy

Cells were imaged using a spinning disk confocal microscope (Carl Zeiss MicroImaging, Thornwood, NY, USA), equipped with a CSU-X1A 5000 spinning disk unit (Yokogawa Electric Corporation, Japan), multi laser module with wavelengths of 458 nm, 488 nm, 514 nm, 561 nm, and 647 nm, and an Axio Observer Z1 motorized inverted microscope equipped with a precision motorized XY stage (Zeiss). Images were acquired with a Zeiss Plan-Neofluar 40× 1.3 NA or 63×1.4 NA objective on an Orca R2 CCD camera using Zen 2.5 software (Zeiss). HeLa cells were plated on dishes containing glass coverslips coated with fibronectin (Mattek Corp. Ashland, MA, USA) 24 h prior to treatment. For time-lapse experiments, OCI-AML cells in media supplemented with HEPES (20 mM) and 2-mercaptoethanol (55 µM) were loaded onto an ethylcellulose micro-scaffold to restrict movement. Cells were allowed to equilibrate to prior to focusing on the cells in a humidified incubation chamber set at 37°C with 5% CO_2_.

### Micro-scaffold preparation and assembly

The micro-scaffold was fabricated as a thin film by soft-lithography. Briefly, ethylcellulose solution (Sigma Aldrich; 7.5% w/v in ethanol) was transferred onto a template containing an array of square-shaped posts (50 x 50 µm; 30 µm height) and let dry at 70 °C for 3 hrs. The micro-scaffold containing square-shaped wells was gently peeled away from the template and kept in a sealed container. The integrity of the wells was confirmed by light microscope. The internal glass surface of 35 mm glass bottom dishes was cleaned using a Q-tip cotton swab dipped in acetone and briefly washed three times with Milli-Q water. Next, a solution of 1M NaOH was incubated overnight before being finally rinsed off three times with Milli-Q water and allowed to dry under a biosafety cabinet airflow. A PDMS solution (SYLGARD 184 Silicone, Dow Corning) was prepared using the manufacturer’s instructions. 10 µL was deposited in the center of the glass bottom dish and spread into a thin film. A square of the micro-scaffold was applied to the PDMS while carefully avoiding trapping air bubbles underneath, then incubated at 37 °C overnight or until the PDMS was fully polymerized. The dishes were UV sterilized by a handheld germicidal UV light (Genesee), followed by sonication in sterile water for 5 min to dislodge air bubbles. The dishes were washed with sterile PBS, prior to seeding the cells.

### Image stream analysis

OCI-AML2 and OCI-AML3 cells stably expressing the C2-Pro BiFC reporter were analyzed using an Amnis ImageStream Mark II (405 nm, 488 nm, 561 nm, and 642 nm excitation lasers; 40× magnification) and INSPIRE software (Luminex). Using the bright-field channel, a cell area-versus-cell aspect ratio plot was used to gate single cells from beads and cellular debris. Subsequently, out-of-focus objects were gated out from the data set using the gradient root-mean-square (RMS) measurement (<45%) on the same bright-field channel. Non-transduced cells were used as fluorescence-minus-one (FMO) controls for the gating strategy. Data analysis was performed using IDEAS software (version 6.2).

### Transient transfection and immunofluorescence

1 x 10^5^ HeLa cells plated on dishes containing glass coverslips coated with fibronectin (1 mg/mL). Cells were transfected with relevant expression plasmids as described in the figure legends using Lipofectamine 2000 transfection reagent (Thermo Fisher Scientific) according to manufacturer’s instructions. After 24 h, cells were washed in 3 x 2 mL PBS and fixed in 2% (w/v) paraformaldehyde in PBS pH 7.2 for 15 min. Cells were washed with PBS followed by permeabilization with 0.15% (v/v) Triton X 100 for 15 min. Cells were blocked in 2% (w/v) BSA for 30 min. Cells were stained with relevant antibodies in 1:100 dilutions in PBS with 2% (w/v) BSA for 1 h. After washing in PBS with 2% (w/v) BSA, the cells were incubated with AF 488 or AF647 conjugated secondary antibodies (Thermo Fisher Scientific) at a 1:250 dilutions in 2% (w/v) BSA for 45 min. Cells were washed in PBS prior to imaging.

### Viability assay

1 X10^4^ parental and caspase-2 deficient *NPM1wt* and *NPM1c+* cells were plated per well in 96 well plates. The plates were incubated for 1, 2, 3, 4 and 5 days at 37⁰C. After each respected day of incubation, the cells were gently centrifuged at 1000 rpm for 2 min to collect the suspension cells. The medium from each well was removed and replaced with fresh medium containing 10 µl of 12 mM MTT as per the manufacturer’s instruction (Roche). After 4 h of incubation, MTT medium was replaced by 100 µL of DMSO followed by orbital plate shaking. The plates were then incubated for an additional 15 min. The absorbance was read at 540 and 680 nm wavelengths.

### RNAseq analysis

Total RNA was extracted from the cells using RNAeasy kit with QIAshredder (Qiagen) according to the manufacturer’s protocol. RNA was resuspended in diethylpyrocarbonate (DEPC) treated water and quantified using a NanoDrop 2000 spectrophotometer (Thermo Fisher Scientific). Quality testing and RNA-seq was carried out at the BGI Americas facility (San Jose, CA). RNA-seq libraries were sequenced using the BGISEQ-500 platform using the DNA nanoball technology (https://completegenomics.mgiamericas.com/en/technology (Shenzhen, China)). Base-calling was performed by the BGISEQ-500 software version 0.3.8.1111. Bioinformatics analysis was carried out using Dr.Tom, (BGI) and ENRICHR pathway analysis tools (https://maayanlab.cloud/Enrichr/). Pathway analysis was carried out on the complete data set (13,043 genes were included for analysis). Significantly differentially expressed genes (DEGs) shared between both caspase-2 deficient clones (813) genes were selected to generate the heat map. The heatmap was generated using R version 4.3.0 and RStudio version 2022.12.0.

### Reverse phase protein array (RPPA) analysis

Reverse phase protein array assays for antibodies to proteins or phosphorylated proteins in different functional pathways were carried out as described previously (Coarfa et al., 2021; Wang et al., 2022b). Specifically, protein lysates were prepared from cultured cells with modified Tissue Protein Extraction Reagent (TPER) (Thermo Fisher Scientific) and a cocktail of protease and phosphatase inhibitors (Roche, Pleasanton, CA). Three technical replicates were spotted for each of the four independent biological replicates for each cell line. The lysates were diluted into 0.5 mg/mL in SDS sample buffer and denatured on the same day. The Quanterix 2470 Arrayer (Quanterix, Billerica, MA) with a 40 pin (185 µm) configuration was used to spot samples and control lysates onto nitrocellulose-coated slides (Grace Bio-labs, Bend, OR) using an array format of 960 lysates/slide (2880 spots/slide). The slides were processed as described and probed with a set of 258 antibodies against total proteins and phosphoproteins using an automated slide stainer Autolink 48 (Dako, Santa Clara, CA). Each slide was incubated with one specific primary antibody and a negative control slide was incubated with antibody diluent without any primary antibody. Primary antibody binding was detected using a biotinylated secondary antibody followed by streptavidin-conjugated IRDye680 fluorophore (LI-COR Biosciences, Lincoln, NE). Total protein content of each spotted lysate was assessed by fluorescent staining with Sypro Ruby Protein Blot Stain according to the manufacturer’s instructions (Molecular Probes, Eugene, OR). Fluorescence-labeled slides were scanned on a GenePix 4400 AL scanner, along with accompanying negative control slides, at an appropriate PMT to obtain optimal signal for this specific set of samples. The images were analyzed with GenePix Pro 7.0 (Molecular Devices, Silicon Valley, CA). Total fluorescence signal intensities of each spot were obtained after subtraction of the local background signal for each slide and were then normalized for variation in total protein, background and non-specific labeling using a group-based normalization method as described (Wang et al., 2022b). For each spot on the array, the-background-subtracted foreground signal intensity was subtracted by the corresponding signal intensity of the negative control slide (omission of primary antibody) and then normalized to the corresponding signal intensity of total protein for that spot. Each image, along with its normalized data, was evaluated for quality through manual inspection and control samples. Antibody slides that failed the quality inspection were either repeated at the end of the staining runs or removed before data reporting. A total of 258 antibodies remained in the list. Antibodies were filtered to remove samples with > 25% CV across the technical replicates and less than 200 signal. A total of 208 antibodies were used for subsequent data analyses. Multiple t-tests were used to determine significance with a cut off of q < 0.01. Z-prime scores and heat maps were generated using Heatmapper (http://www.heatmapper.ca).

### Statistical analysis

Statistical significance was assessed by using two-tailed Student’s t test. Two-way ANOVA was used to correct for multiple comparisons. All statistical analyses were performed using GraphPad Prism 9 for Windows (San Diego, CA, and USA RRID: SCR_002798).

## Acknowledgments

We would like to thank Suruchi Salgar for computational assistance and the entire Bouchier-Hayes lab for helpful discussion. Funding for this project includes NIH/NIGMS R01GM121389 (LBH), NIH/NCI R21CA256606 (LBH) and CPRIT RP210027. We would like to acknowledge the Texas Children’s Hospital William T. Shearer Center for Human Immunobiology for their generous support for this research and the expert assistance of Daniela Carrasco Di Lallo and Dr. Alexander Vargas-Hernandez. This work was supported in part by Cancer Prevention & Research Institute of Texas Proteomics & Metabolomics Core Facility Support Award (RP210227) and NCI Cancer Center Support Grant (P30CA125123) to the Antibody-based Proteomics Core/Shared Resource (SH) and NIH S10 award (S10OD028648, SH). We thank Drs. Xuan Wang and Zhongcheng Shi from the Antibody-based Proteomics Core/Shared Resource for their excellent technical assistant in performing RPPA experiments. We thank Drs. Cristian Coarfa and Sandra L. Grimm for RPPA data processing and normalization. Graphics were created using BioRender.

