## Supplemental Text for "Caspase-2 is essential for proliferation and self-renewal of nucleophosmin-mutated acute myeloid leukemia"

***Supplemental Movies***

**Supplemental Movie S1.** **Daunorubicin-induced caspase-2 BiFC in *NPM1wt* cells.** Movie of representative OCI-AML2 (*NPM1wt)*.C2 Pro-BiFC cells treated with daunorubicin (0.5 µM).  Images were acquired by confocal microscopy every 10 min for 16 h with the appropriate setting. The movie shows cells expressing mCherry throughout the cell (*red*) and caspase-2 BiFC (*yellow*) in the nucleolus and cytoplasm. Bar, 10 µm.

**Supplemental Movie S2.** **Daunorubicin-induced caspase-2 BiFC in *NPM1c+* cells.** Movie of representative OCI-AML3(*NPM1c+*).C2 Pro-BiFC cells treated with daunorubicin (0.5 µM).  Images were acquired by confocal microscopy every 10 min for 16 h with the appropriate setting. The movie shows cells expressing mCherry throughout the cell (*red*) and caspase-2 BiFC (*yellow*) in the nucleolus and cytoplasm. Bar, 10 µm.
