## Supplemental figures for "Caspase-2 is essential for proliferation and self-renewal of nucleophosmin-mutated acute myeloid leukemia"

Figure S1

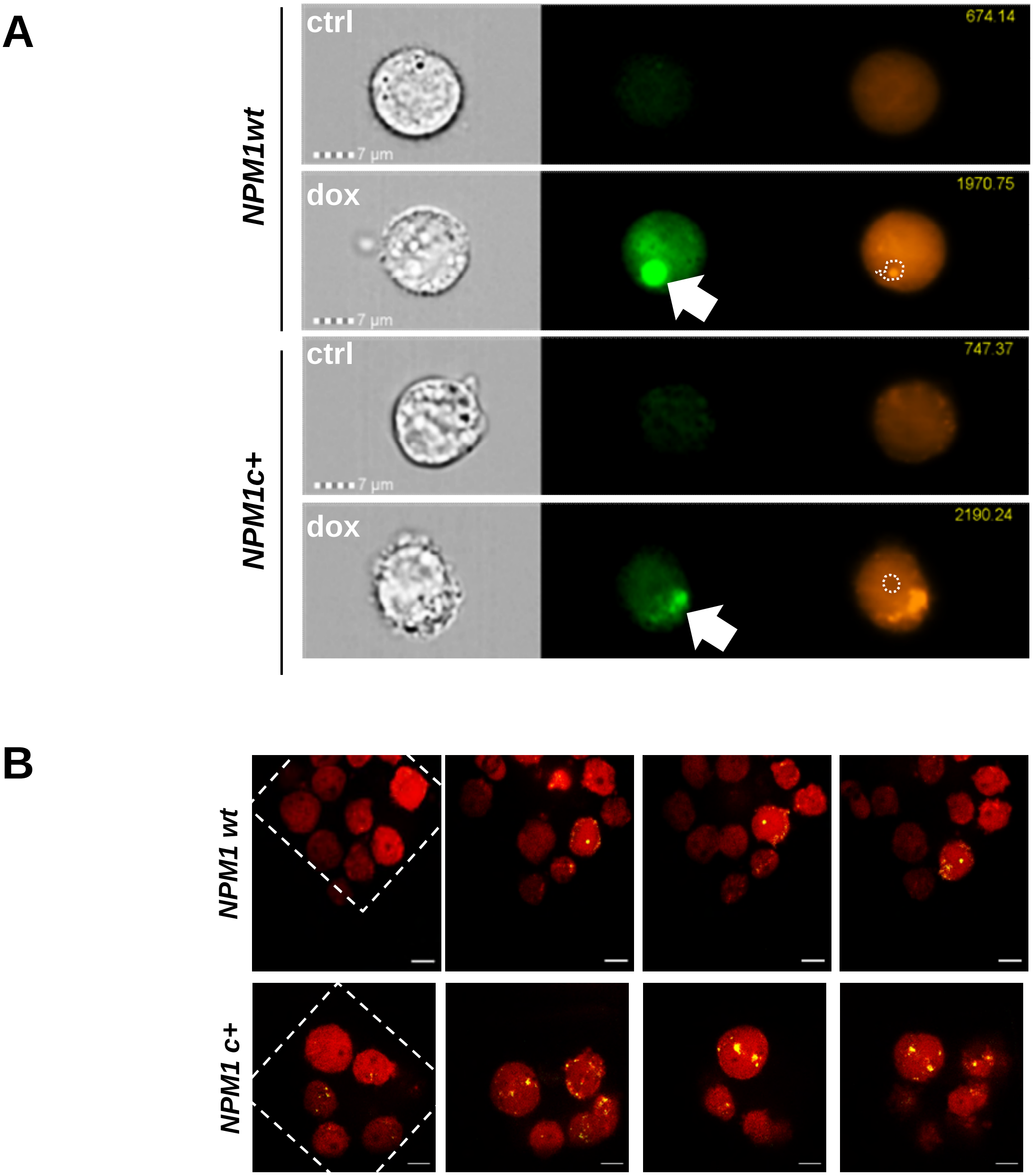


**Figure S1: Caspase-2 is activated in the same compartment as NPM1. (A)** *NPM1wt* and *NPM1c+* cells stably expressing C2-Pro VC-2A-C2-Pro VN-2A-mCherry (C2-Pro BiFC) were treated with doxorubicin (dox, 0.5 µg/ml) for 16 h in the presence of the caspase inhibitor qVD-OPH (5 µM) to prevent cell death caused by apoptosis. Representative images of cells acquired with an imaging flow cytometer show total cell volume (mCherry, orange) and caspase-2 BiFC signal (green, arrows). The dotted outline shows the position of the nucleolus. (B) *NPM1wt* and *NPM1c+* C2-Pro BiFC cells were plated on a micro-scaffold comprised of 50 $\mu m^2$ squares, treated with daunorubicin (0.5 µM) , and imaged by confocal microscopy every 5 min for 16 h. The white dotted line delineates the grid enabling stable positioning of the cells. Yellow =Venus (Caspase-2 BiFC); red=mcherry. Representative images of time-lapses are shown. Bar, 20 µm.

Figure S2

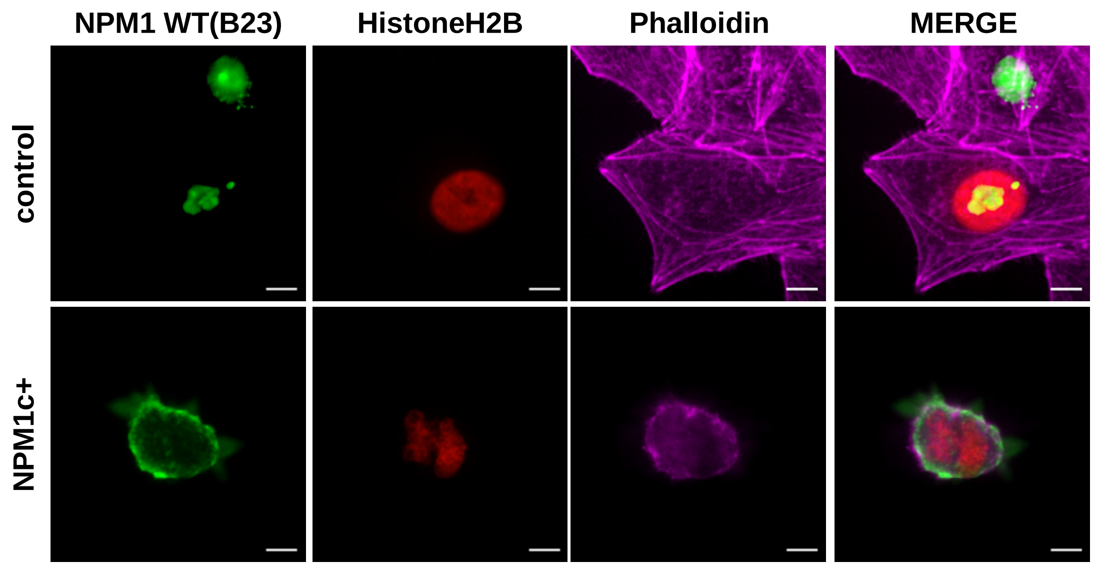


**Figure S2: NPM1c+ relocalizes NPM1 to the cytoplasm** (A) HeLa cells were transiently transfected with an empty vector or an expression construct for *NPM1c+* along with H2B mCherry as transfection reporter. Cells were stained with phalloidin to show the actin cytoskeleton in the cytoplasm (purple), and anti-B23 (green) that recognizes only wild-type NPM1. H2B-mCherry delineates the nucleus (red), Representative images taken 24 h post transfection are shown. Bar, 10 µm.

Figure S3

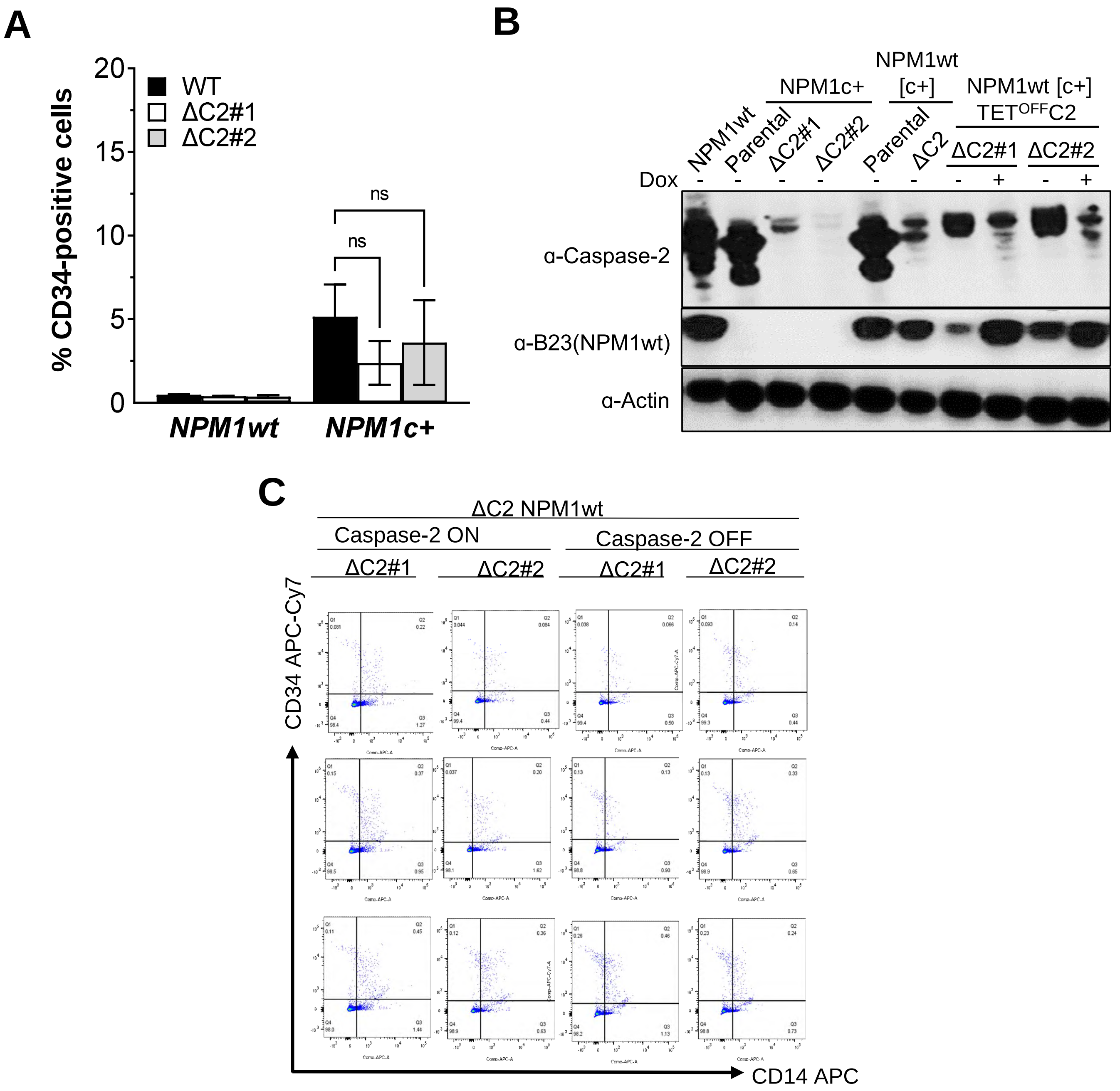


**Figure S3: Loss of caspase-2 induces NPM1c+ dependent differentiation.** (A) Cell surface expression of CD34 was measured in OCI-AML-2 (*NPM1wt*) and OCI-AML-3 (*NPM1c+*) parental or ΔC2 cells cultured for two weeks by flow cytometry. The percentage of cells expressing CD34 is shown. Results are the average of three independent experiments plus or minus standard deviation. (B) Lysates from the indicated cell lines treated with or without doxycycline (2 µg/mL) for 16 h to repress caspase-2 expression were immunoblotted for caspase-2, wild-type NPM1 (B-23), and actin as a loading control. (C) *NPM1wt*C2$^{OFF}$ cells were grown in DMSO (C2ON) or doxycycline (2 µg/mL, C2OFF) for 3 weeks. Cell surface CD14 and CD34 expression was measured once a week by flow cytometry. Representative flow plots are shown.

Figure S4

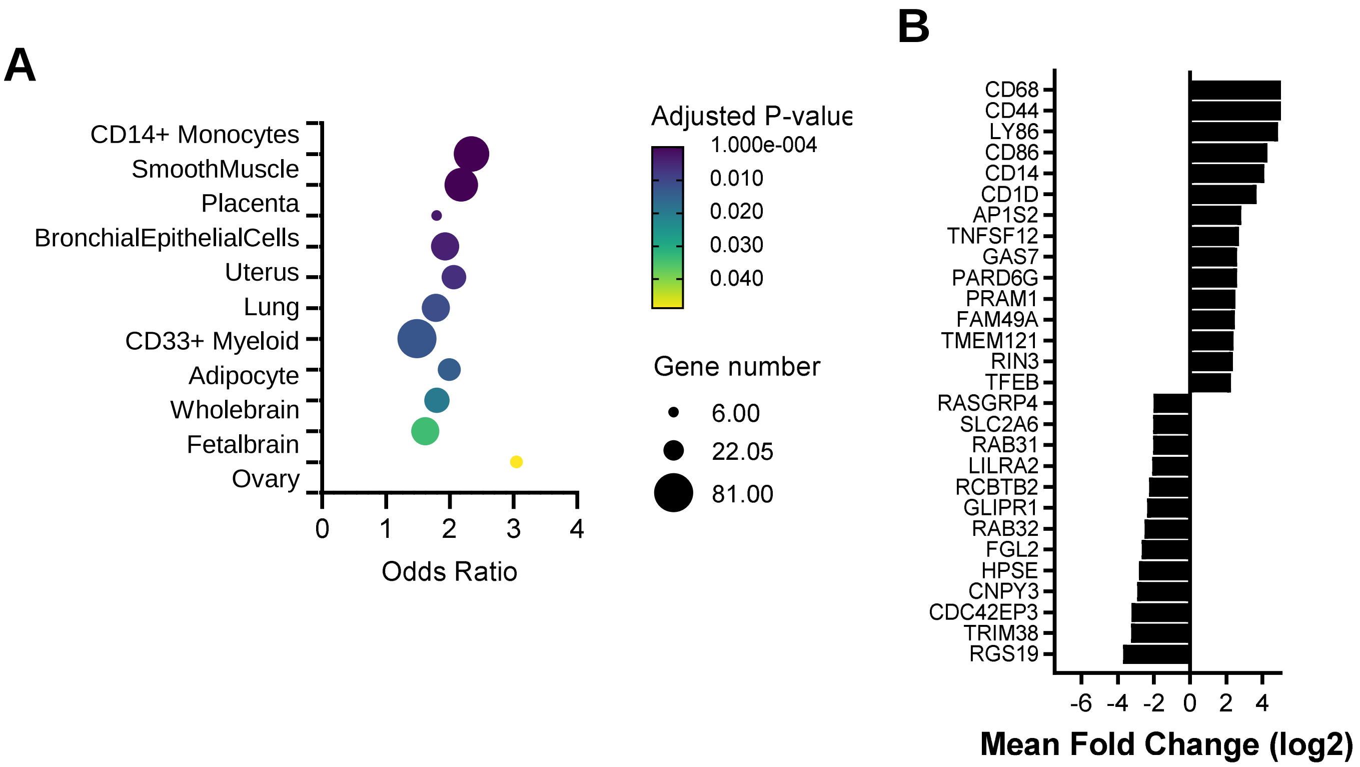


**Figure S4: Cell type changes in NPM1c+ versus. caspase-2 knockout cells. (**A) Total RNAseq data was analyzed with ENRICHR to differentially regulated genes between *NPM1c+* parental and ΔC2 cells associated with the indicated cell types. The size of the dots represent the number of genes associated with each biological pathway. The color of the dots represents the significance (adjusted p-value). B) The mean log2 fold change of significant DEGs between NPM1c+ parental and ΔC2 cells involved in the KEGG term CD14+ monocytes is shown.
